# GSK-3β coordinates axonal microtubule organisation through Shot and Tau

**DOI:** 10.1101/2024.09.08.611864

**Authors:** Andre Voelzmann, Lubna Nuhu-Soso, Alex Roof, Sanjai Patel, Hayley Bennett, Antony Adamson, Marvin Bentley, Gareth J. O. Evans, Ines Hahn

## Abstract

Glycogen Synthase Kinase 3β (GSK-3β) is a key coordinator of neuronal development and maintenance; hyperactive GSK-3β is linked to neurodevelopmental and -degenerative diseases and therefore a promising therapeutic target. In neurons, GSK-3β coordinates the cytoskeleton by phosphorylating microtubule-binding proteins. In this study, we found that tight regulation of GSK-3β kinase activity is required for the maintenance of parallel microtubule bundles in *Drosophila* and rat axons. Up- or down-regulation of GSK-3β led to axons forming pathological swellings in which microtubule bundles disintegrated into disorganised, curled microtubules. We identified the microtubule bundling proteins Shot and Tau as key GSK-3β targets and found that GSK-3β exerted its regulatory effect on microtubule bundling through them. GSK-3β regulates the ability of Shot and Tau to attach to microtubules and/or the plus-end protein Eb1. Mis-regulation of GSK-3β leads to the loss of Eb1–Shot-mediated guidance of polymerising microtubules into parallel bundles, thus causing disorganisation. We propose microtubule disorganisation as a new explanation for how GSK-3β hyperactivity leads to neurodegeneration and why global inhibition of GSK-3β has not been successful in clinical trials for neuronal disorders.

## Introduction

The development and maintenance of axons critically depend on parallel microtubule bundles. Microtubules establish axonal projections, adapt neuronal cell shape, and are the substrate for motor proteins that move vesicles and other cargoes ^1–3^. The regulation of microtubule polymerisation and stability is crucial for the formation of microtubule bundles, and disruptions to this process—such as microtubule loss or disorganization—are common features in many axon pathologies ^3,4^. Despite their physiological importance, the molecular mechanisms that maintain microtubule bundles are not understood ^5^.

The kinase GSK-3β has emerged as a key regulator of microtubule stability and dynamics ^6^. GSK-3β is required for proper neuronal development and mis-regulation of GSK-3β is linked to several neurodevelopmental and -degenerative disorders ^7–11^. GSK-3β is therefore a promising therapeutic target for neurological disorders. However, global GSK-3β inhibition has either not been therapeutically beneficial ^12^ or lead to long-term toxicity (e.g. approved inhibitor lithium) ^13^ and neuronal decay ^14^, indicating a not yet appreciated complexity of GSK-3β regulation.

Here we tested if mis-regulation of GSK-3β affects axonal microtubule bundles in primary neuron cultures from genetically tractable *Drosophila* ^5,15–19^ and primary rat hippocampal neurons. We found that mis-regulation of GSK-3β—both hyper- and in-activation—disrupted the organisation of parallel microtubule bundles. We identified two key microtubule-associated proteins, the spectraplakin Short Stop (Shot) and Tau, as essential GSK-3β targets for proper microtubule bundling. Both proteins play an important role in maintaining microtubule bundles^16,18^.

We found that hyperactivation of GSK-3β causes microtubule unbundling by reducing binding of Tau to microtubules, which disrupts the Shot/Eb1-mediated alignment of polymerizing microtubules into parallel bundles ^18^. In contrast, GSK-3β inactivation detaches the microtubule-actin crosslinker Shot from plus ends. This prevents Shot from guiding polymerizing microtubules into proper bundles. These differential effects of GSK-3β mediated modulation of key microtubules regulators and their impact on microtubule bundle maintenance is a new model to explain how hyper- and hypo-phosphorylation by GSK-3β causes pathology. Furthermore, this framework explains why global GSK-3β inhibition has yielded little to no therapeutic effect on neurodegenerative diseases.

## Methods

### Fly stocks

Lines for targeted gene expression were *UAS-sgg.S9A* (*UAS-sggCA*; constitutively active (CA) form of Sgg, ^20^)*, UAS-sgg.A81T (UAS-sgg.DN;* inactive form of Sgg, ^21^), *UAS-Eb1-GFP and UAS-shot-Ctail-GFP (EGC-GFP)*^16^. Loss-of-function mutant stocks used in this study were *sgg*^1^ (hypermorphic allele, ^22^), *dtau^KO^* (a null allele; ^23^, tau-/-), *shot*^3^ (the strongest available allele of *short stop*; ^24,25^). Gal4 driver lines used were the pan-neuronal lines *elav-Gal4* (1^st^ and 3^rd^ chromosomal, both expressing at all stages) ^26^. Protein trap line used was P{Wee}tau[304] (tauGFP; ^27,28^). To induce gene expression with RU486, UAS constructs were expressed using an elavGal4-GeneSwitch driver line (RRID:BDSC_43642, ^29,30^). Gene expression was induced by adding 200 mg/mL RU486 to cell culture media. Oregon R stocks, heterozygous crosses of UAS or Gal4 lines with Oregon R or uninduced fly lines were used as controls as indicated.

### Generation of Shot-eGFP CRISPR/Cas9 lines

The five C-terminal exons of Shot coding for the C-tail and SxIP sites (shot-RH, FBtr0087621) were excised from the shot genomic region and replaced with fused version of those exons and an additional C-terminal eGFP or a phosphodeficient/mimetic version following the same strategy, leading to either eGFP-tagged wildtype Shot (*Shot^WT^-eGFP*) containing a unmodified GSK-3β target site (SRAGSKPNSRPLSRQGSKPPSRHGS), phospho-deficient Shot where serines where mutated into alanines (**A**RAG**A**KPN**A**RPL**A**RQG**A**KPP**A**RHG**A**), or phospho-mimetic Shot where serines were mutated into aspartic acid residues (**D**RAG**D**KPN**D**RPL**D**RQG**D**KPP**D**RHG**D**). The three Shot coding sequences with eGFP were synthesised as gBlocks double stranded DNA fragments (IDT).

Suitable gRNA target sites (5′ gRNA: GGAGGCTCTCGTGCCGGCTC, 3′ gRNA: GCTATAGGAAGCCACCGTTA) were identified by CRISPR optimal target finder ^31^ and cloned into pCFD5 (Addgene plasmid #73914; ^32^) via Gibson assembly (NEB).

CRISPR donor plasmids were cloned with 0.8 kb 5’ and 3’ homology arms amplified from fly genomic DNA into pUC57(Addgene plasmid #51019, RRID:Addgene_51019), flanking the different CTail Shot-eGFP coding sequences, amplified from the above gBlocks and assembled using a three-fragment Gibson reaction. Primer sequences are provided below.

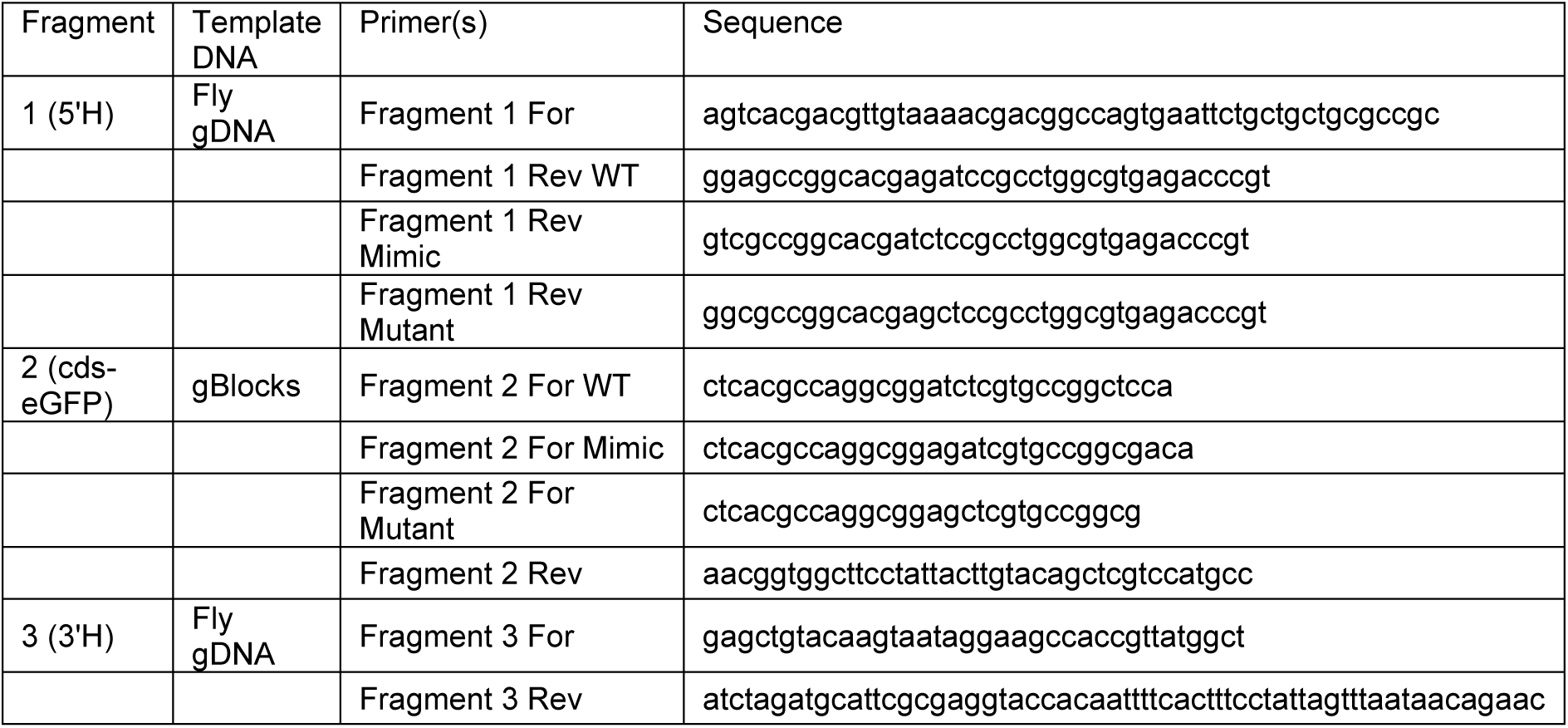

Constructs were midi-prepped (Bioline) and eluted in ddH_2_O, before injection into y[1] sc[1] v[1];; {y[+t7.7] v[+t1.8]=nanos-Cas9}attp2 (gift from the University of Cambridge) and selected for eGFP-positive flies. Positive candidates were confirmed by Sanger sequencing.

### *Drosophila* primary cell culture

*Drosophila* primary neuron cultures followed protocols described previously ^17,18,33^: Embryos (stage 11) were incubated with bleach for 90s, sterilized in 70% ethanol (not exceeding 5 min), washed in sterile Schneider’s medium supplemented with 20% fetal calf serum (Schneider’s/FCS; Gibco), and homogenized with micro-pestles in 100 μl dispersion medium (0.005% phenylthiourea, 1% Penicilin/Streptomycin in HBSS medium (SigmaAldrich, ^33^), and incubated for 4 min at 37°C. Dispersion was stopped with 200 μl Schneider’s/FCS, cells were spun down for 4 min at 650 g, supernatant was removed and cells re-suspended in 30 µl of Schneider’s/FCS per cover slip. Usually, each of the >three biological repeats consisted of three coverslips (technical repeats) were cultured for each condition with 6-8 embryos per coverslip (18-24 for three). 30 μl drops were placed in the centre of culture chambers and covered with a concanavalin A (5 µg/ml) coated coverslip. Cells were allowed to adhere to coverslips for 90-120 min, then turned and grown as a hanging drop culture at 26°C as indicated.

### Rat hippocampal primary cell culture

Primary hippocampal neurons were cultured as previously described ^34^. In short, E18 rat hippocampi were dissected, trypsinised, dissociated, and plated on 18 mm glass coverslips coated with poly-l-lysine. Cultured neurons were grown in N2-supplemented Minimal Essential Medium at 37°C with 5% CO2. Stage 4/5 hippocampal neurons (5-11 DIV respectively) were transfected with Lipofectamine 2000 (Thermo Fisher, Cat# 11668019) following manufacturer’s instructions. To image stage 3 neurons, dissociated neurons were electroporated by nucleofection (AMAXA) using standard protocols prior to plating. Cells were allowed to express for 24 h before imaging or fixation.

### Immunohistochemistry

Primary neurons were fixed for 30 min at room temperature (4% paraformaldehyde (PFA) in 0.05 M phosphate buffer, pH 7–7.2). For anti-Eb1 staining ice-cold +TIP fix (90% methanol, 3% formaldehyde, 5 mM sodium carbonate, pH 9; stored at -80°C and added to the cells) ^35^ was added and cells were fixed for 10 mins at -20°C and then washed in PBT (PBS with 0.3% TritonX). All subsequent staining and washing steps were performed with PBT.

The following staining reagents were used: anti-tubulin (clone DM1A, mouse, 1:1000, Sigma); anti-DmEb1 (gift from H. Ohkura; rabbit, 1:2000) ^36^; anti-GFP (rab, 1:500, ab290, abcam); anti-HA (rat, 1:200, 3F10, SigmaAldrich); F-actin was stained with Phalloidin conjugated with TRITC/Alexa647, FITC or Atto647N (1:100; Invitrogen and Sigma). Specimens were embedded in ProLong Gold Antifade Mountant (ThermoFisher Scientific).

### Microscopy and data analysis

For standard imaging we used AxioCam 506 monochrome (Carl Zeiss Ltd.) or MatrixVision mvBlueFox3-M2 2124G digital cameras mounted on BX50WI or BX51 Olympus compound fluorescence microscopes. For the analysis of primary neurons, we used the following parameters:

Degree of disorganised microtubule curling in axons was assessed as “microtubule disorganisation index” (MDI; ^17,18^): Axon length is marked using the segmented line tool of Image J (we measured length from cell body to growth cone tip ^18,37^) and the area of disorganised microtubules was measured with the freehand selection tool in ImageJ. The MDI was calculated by dividing the sum of areas of disorganisaiton by axon length. This process was automated using the axondisorg_table (https://github.com/avmann/axondisorg_table) or axon-disorganisation-from-rois (https://github.com/avmann/axon-disorganisation-from-rois) Fiji/ImageJ macros.

The amount of Eb1 at microtubule comets was approximated from Eb1 staining on microtubule ends by multiplying comet mean intensity with comet length (described here ^18^).

### Recombinant protein expression and purification

Primers were designed for MAP1B, shot and shot mutant peptide sequences with 5’ EcoRI and 3’ XhoI sites, annealed and ligated into pGEX-4T-1 (GE Healthcare). Peptide sequences are as follows; ERLSPAKSPSLSPSPPSPIEKT for MAP1B, SRAGSKPNSRPLSRQGSKPPSRHGS for shot and ARAGAKPNARPLARQGAKPPARHGA for shot mutant. Empty pGEX-4T-1 plasmid was used for GST only.

All constructs were expressed in BL21 E. coli at 37°C. Cultures at OD600 = 0.8 were induced with 1 mM Isopropyl β-D-1-thiogalactopyranoside (IPTG) for 3 h at 37°C with shaking at 200 rpm. Bacteria were harvested by centrifugation at 5000 x g for 15 min at 4°C and pellets were stored at -70°C overnight. Pellets were resuspended in 20 ml of PBS containing 1 mM phenylmethylsulfonyl fluoride (PMSF), 1X protease inhibitor cocktail and 6.65 mg of lysozyme. After incubation on ice for 30 min, 1 % (v/v) Triton X-100 and 5 mM DTT were added followed by 6 x sonication on ice for 30 s on and off for a total of 5 min 30 s at 17 kHz. The lysate was clarified by centrifugation at 12,000 rpm for 30 min at 4°C and the supernatant was incubated with 3x PBS pre-washed Glutathione Sepharose 4B resin for at least 1 h at 4°C with agitation. Glutathione resin was recovered by centrifugation, washed and protein was eluted in elution buffer (100 mM Tris pH 8.0, 20 mM glutathione, 100 mM NaCl). Purified protein was stored at - 70°C until use.

### In vitro kinase assays

Constitutively active GSK-3β protein was obtained from MRC PPU (Dundee). Rabbit monoclonal thiophosphate ester antibody (clone 51-8) and p-Nitrobenzyl mesylate (PNBM) were obtained from Abcam (Cambridge, UK). ATPγS was obtained from Biorbyt (Cambridge, UK). All other reagents were obtained from Sigma (Dorset, UK)

Thiophosphorylation assays were performed in 25 µl reaction volumes at 30°C for 3 h. The reactions contained 400 ng of GSK-3β, 3 µg of substrate (GST or GST-peptides) and reaction buffer (50 mM HEPES pH 7.5, 10 mM MgCl2, 100 mM NaCl, 1 mM DTT). The reactions were left to equilibrate at 30°C after which ATPγS was added to a final concentration of 1 mM to start the reaction. After 3 h, the alkylating agent, PNBM, was added to a final concentration of 2.5 mM and reactions were left for 2 h at room temperature. Reactions were quenched by addition of 4X Laemmli sample buffer. Samples were separated on a 12.5% SDS-PAGE gel, transferred to PVDF membrane, and thiophosphorylation determined by Western blotting with rabbit anti-thiophosphate ester (1:10,000) and anti-rabbit-HRP (1:5,000) and visualised by enhanced chemiluminescence reagent (ECL). Coomassie-stained gels were run alongside to ensure equal loading of substrates. Densitometry was performed on Western blots and Coomassie-stained gels using ImageJ.

## Results

### Activation or inactivation of GSK-3β cause disorganisation of axonal microtubules in *Drosophila* primary neurons

To evaluate a role of GSK-3β in axonal microtubule organisation, we expressed constitutively active or dominant-negative GSK-3β mutants in cultured neurons and analysed microtubule organisation. Primary neurons were derived from embryos of fly lines overexpressing either constitutively active GSK-3β (CA, *UAS-sggCA*, S9A mutation ^21^) or dominant-negative, kinase-dead (*UAS-sggDN*; A81T mutation ^21^). After 6 h in vitro (HIV), neurons were fixed and immuno-stained for tubulin to visualize microtubules. Activation or inhibition of GSK-3β caused axon swellings. Within these swellings microtubules lost their bundled conformation and became curly (**Fig. 1A**). We quantified this phenotype using the previously described microtubule disorganisation index (MDI; area of microtubule disorganisation relative to axon length; see ^17,18,38,39^). The MDI measures the fold-increase of microtubule unbundling relative to control neurons. The measurements resulted in an MDI of 1.8 for *sggCA* and 2.1 for *sggDN* (**Fig. 1C**). To independently verify the impact of GSK-3β inactivation on microtubule organisation, we cultured neurons from animals with a GSK-3β loss-of-function allele (*sgg*^1^ ^40^) or treated wild type neurons with increasing concentrations of the GSK-3β inhibitor lithium (LiCl ^41^). Both treatments resulted in a significant increase in microtubule unbundling (*sgg*^1^ MDI=1.8; 10 mM lithium MDI=3.3) (**Fig. 1D**). To determine if GSK-3β regulation of microtubule organisation is limited to early development or required in all developmental stages, we analysed neurons expressing *sggCA* or *sggDN* for three days in vitro (3 DIV, **Fig. 1B**). Older neurons also exhibited increased microtubule curling in both conditions (*sggCA* MDI=2.5; *sggDN* MDI=2.7) (**Fig. 1C**). Furthermore, we observed significant microtubule unbundling in neurons cultured from larval brains expressing *sggCA* (MDI=3.9) or *sggDN* (MDI=2.8) (**Fig. 1D**). Because both over- and inactivation of GSK-3β resulted in microtubule curling, GSK-3β activity must be tightly regulated in all stages of neuronal development to efficiently maintain microtubule bundles.

**Figure 1.**
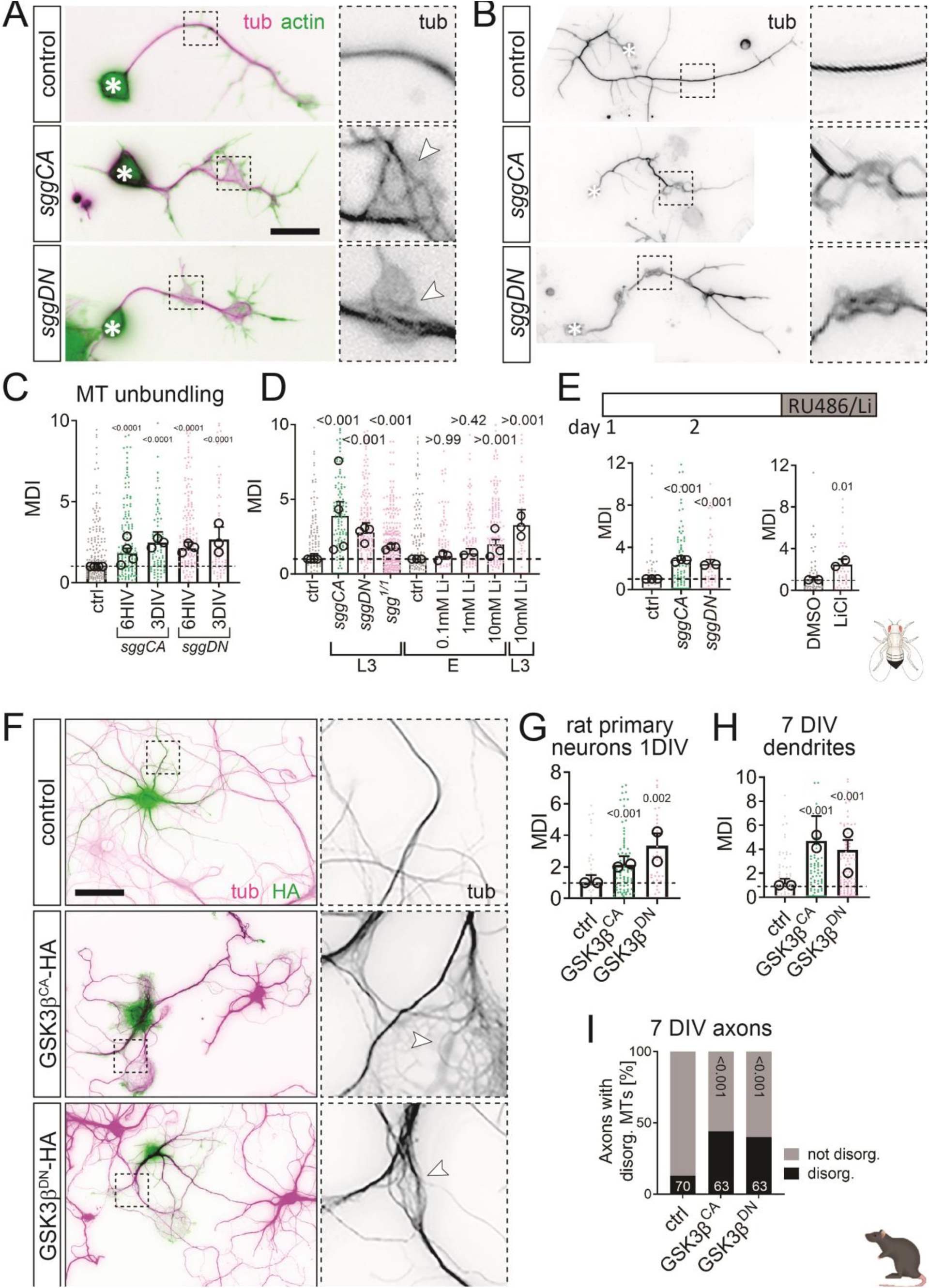
Inhibition and hyper-activation of GSK-3β/Shaggy both lead to curling and unbundling of axonal microtubules in *Drosophila* primary and rat hippocampal neurons. **A, B**) Images of representative examples of *Drosophila* embryonic primary neurons either immuno-stained for tubulin and actin (A) or tubulin (B). Neurons of the following conditions were cultured for 6 hours in vitro (6HIV, A) or 3 days in vitro (3DIV, B): controls (ctrl; elavGal4, UAS-GFP), expressing constitutively active (*sggCA*; *elavGal4, UAS-sgg^S9A^*) or inactive (*sggDN*; *elavGal4, UAS-sgg^A81T^*) UAS-GSK-3β variants via the pan-neuronal driver elavGal4. Asterisks indicate cell bodies, dashed squares in are shown as 3.5-fold magnified close-ups beside each image white arrow heads point at areas of microtubule curling. **C)** Quantification of microtubule unbundling depicted as MDI (see methods) obtained from embryonic primary neurons with the same genotypes as shown in A,B. **D)** Microtubule unbundling quantification of primary neurons obtained from third instar larval brains (L3) expressing in/active GSK-3β (*sggCA/DN*), embryonic primary neurons of GSK-3β mutants (*sgg*^1^*^/^*^1^) and neurons treated with the GSK-3β inhibitor Lithium. **E)** *sggDN* or *sggCA* expression was induced via RU486 (elav::switchGal4) at 3DIV for 1DIV or control neurons were treated with 10mM LiCl. **F)** Images of representative examples of rat hippocampal neurons at 7 DIV expressing HA (controls), GSK-3β^CA^-HA or GSK-3β^DN^-HA immuno-stained for tubulin and HA. **G-I)** Quantification of microtubule unbundling depicted as MDI (G,H) or ratio (axons w/ or w/o microtubule unbundling, I) at conditions indicated above. Data were normalised to parallel controls (dashed horizontal line) and are shown as mean ± SEM; data points in each plot, taken from at least two experimental repeats consisting of 3 replicates each; large open circles in graphs indicate median/mean of independent biological repeats. P-values obtained with Kruskall-Wallis ANOVA test for the different genotypes are indicated in each graph. Scale bar in A represents 10 μm in A and 20 μm in F. For raw data see S1 Data.

We next analysed if altering GSK-3β levels induces microtubule unbundling in mature neurons. We used a modified Gal4/UAS system where transgene expression is induced by the small molecule RU486 ^42^. *ElavGS/UAS-sggCA* or *-sggDN* neurons were grown without induction for three days to mature to the post growth cone stage where they form presynapses^43^. Expression of *sggCA* or *sggDN* was then induced by treatment with 100 µg/ml RU486 for one day. Neurons were fixed, stained for α-tubulin to visualize microtubules, and imaged. Control neurons (i.e., not treated with RU486) exhibited little microtubule disorganisation. In contrast, microtubule disorganisation was significantly increased by expression of constitutively active (*sggCA* MDI=2.7) and dominant negative (*sggDN* MDI=2.3) GSK-3β (**Fig. 1E**).

As an orthogonal approach, we treated three-day old, cultured neurons with the GSK-3β inhibitor Li for 24 h and measured microtubule disorganisation (**Fig. 1E**). 10 mM LiCl treatment resulted in a 2.3-fold increase in microtubule disorganisation relative to vehicle-treated neurons (DMSO), comparable to expression of constitutively active GSK-3β (**Fig. 1E**).

Together, these results indicate that both up- and down-regulation of GSK-3β activity causes microtubule disorganisation. Therefore, GSK-3β activity must be closely regulated for proper microtubule organisation. This regulation of GSK-3β is required during neuron development and axonal outgrowth, and for maintenance of the microtubule network in mature neurons.

### GSK-3β-mediated regulation is required for maintaining microtubule organisation/bundling in mammalian neurons

To explore whether this observation applies universally, we determined if mis-regulation of GSK-3β affects microtubule organisation in cultured hippocampal neurons. We electroporated neurons with constitutively active (HA-GSK-3β^CA^) or dominant-negative (HA-GSK-3β^DN^) GSK-3β plasmids before plating (0 DIV). Neurons were fixed at 1 DIV and stained for α-tubulin and HA. At 1 DIV, most neurons are in developmental stage 2, having extended minor neurites but prior to polarization and the differentiation of axon and dendrites ^44^. After imaging, we generated the MDI for HA-expressing neurons and untransfected neurons (control) on the same coverslips. Expression of constitutively active (MDI=2.1) and dominant negative (MDI=3.3) GSK-3β resulted in a significant increase in microtubule disorganisation (**Fig. 1F,G**). These results show that GSK-3β regulation is required for proper microtubule organisation in developing mammalian neurons.

We then asked if GSK-3β regulation is required for the maintenance of microtubule organisation in mature neurons. We transfected 6 DIV hippocampal neurons with constitutively active or dominant-negative GSK-3β plasmids and fixed 24 h after transfection. At 7 DIV, hippocampal neurons are fully polarized and have developed axons and dendrites ^44^. Both conditions resulted in a significant increase of microtubule disorganisation (**Fig. 1F,H**). Quantifications found that both constructs resulted in a large increase in microtubule disorganisation compared to untransfected neurons (HA-GSK-3β^CA^ MDI=4.6; HA-GSK-3β^DN^ MDI=3.9; **Fig. 1H**). We measured MDI in dendrites and observed a 4.6- and 3.9-fold increase in GSK-3β^CA^ and GSK-3β^DN^ respectively. We were not able to calculate the MDI for axons, because axons of 7 DIV neurons extend substantially beyond the field of view and the entire axon length is a required MDI parameter. Instead, we calculated the percentage of axons with disorganised MTs within each field of view and observed an increase from 13% in untreated neurons to 44% (GSK-3β^CA^) and 40% (GSK-3β^DN^) (**Fig. 1I**).

These experiments show that the effects we saw after modifying GSK-3β activity in *Drosophila* were recapitulated in mammalian neurons. Therefore, the requirement for balanced GSK-3β activity to maintain microtubule bundles is a conserved feature and likely a universal mechanism for maintaining the neuronal cytoskeleton.

### GSK-3β affects Eb1 comet formation differentially

We previously found that microtubule bundle arrangement is tightly linked to the availability of Eb1 at microtubule plus ends, as there is a negative correlation between the size of Eb1 comets and microtubule curling: smaller Eb1 comets result in larger MDI values (**Fig. 2A**) ^18^. We therefore tested if GSK-3β hyper- or in-activation depleted Eb1 at microtubule plus ends.

**Figure 2.**
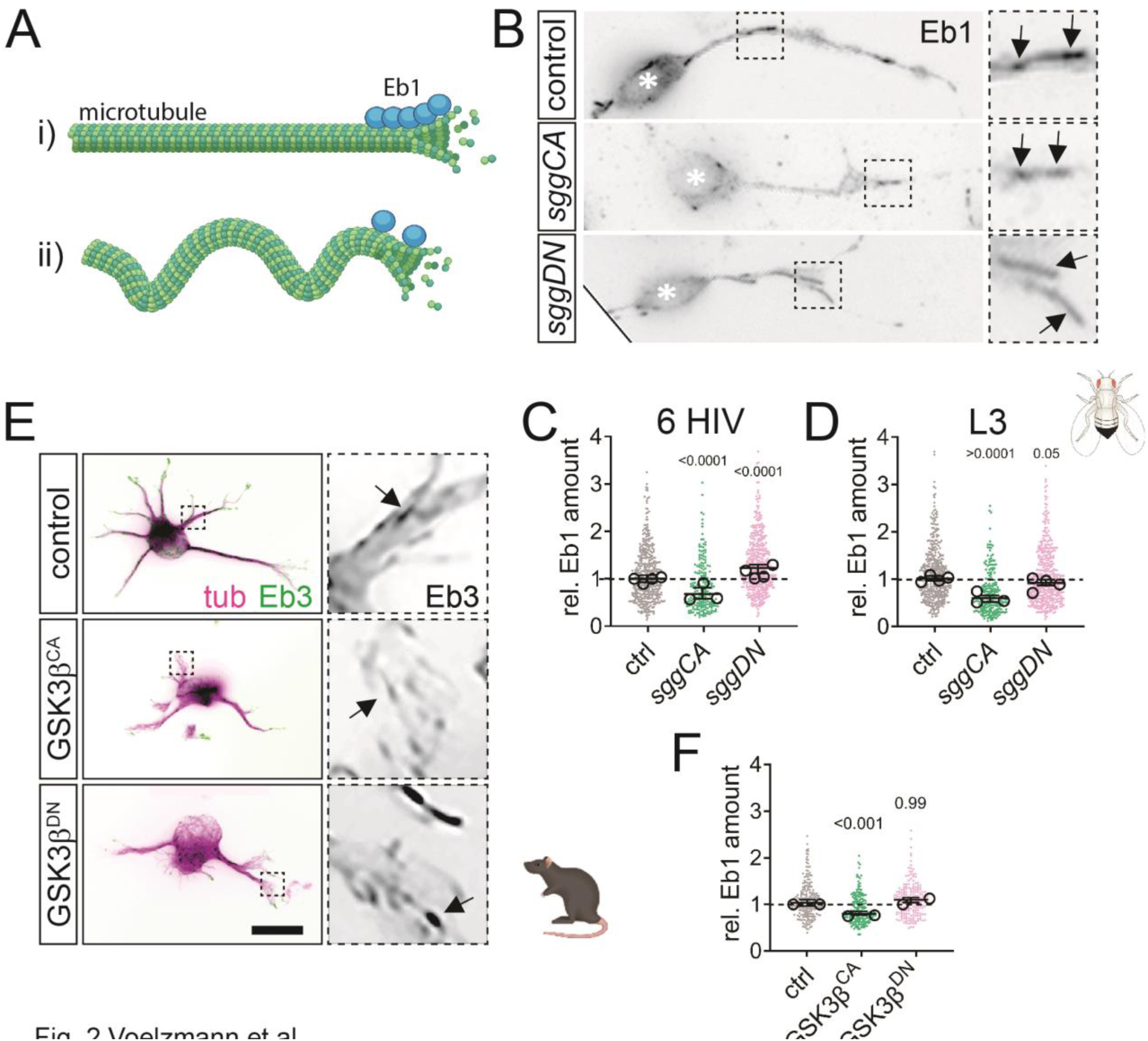
Expression of active GSK-3β reduces Eb1 comet size. **A)** Schematic representation how loss of Eb1 leads to microtubule unbundling (generated with BioRender): i) normal Eb1 levels support microtubule polymerisation and straight microtubule arrangement whereas ii) reduced Eb1 levels lead to microtubule curling. See ^18^ for details. **B)** Images of representative examples of control embryonic primary neurons or neurons expressing constitutively active (*sggCA*) or inactive GSK-3β (*sggDN*) immuno-stained for Eb1. Asterisks indicate cell bodies, dashed squares in are shown as 3.5-fold magnified close-ups beside each image arrows point at Eb1 comets. Scale bar in B represents 10 μm. **C, D)** Quantification of normalised Eb1 amounts at plus ends obtained from embryonic primary neurons at 6HIV and larval neurons after 1 DIV (L3). **E)** Representative images of primary rat hippocampal neurons at 1 DIV co-expressing Eb3-GFP with HA (controls), GSK-3β^CA^-HA or GSK-3β^DN^-HA. Asterisks indicate cell bodies, dashed squares in are shown as 3.5-fold magnified close-ups beside each image arrows point at Eb1 comets **F)** Quantification of normalised Eb1 comet lengths (see Methods) obtained from the same genotypes as shown in E. Data were normalised to parallel controls (dashed horizontal line) and are shown as mean ± SEM; data points in each plot, taken from at least two experimental repeats consisting of 3 replicates each; large open circles in graphs indicate median/mean of independent biological repeats. P-values obtained with Kruskall-Wallis ANOVA test for the different genotypes are indicated in each graph. Scale bar in A represents 10 μm in B and 20 μm in F. For raw data see S2 Data.

We cultured embryonic or larval neurons expressing constitutively active or dominant negative GSK-3β and determined the amount of Eb1 at microtubule plus ends by immuno-staining (**Fig. 2B,C**). We found that Eb1 at microtubule plus ends was decreased by constitutively active GSK3β (*sggCA*) in embryonic (6 HIV, 67%) and larval (L3, 58%) neurons. Expression of dominant-negative GSK-3β (*sggDN*) only mildly affected Eb1 at microtubule plus-ends. Embryonic neurons exhibited more Eb1 at microtubule tips (6 HIV) in contrast to no change of Eb1 at tips of larval neurons (L3). To test if GSK-3β manipulation had comparable effects in mammalian neurons, we analysed Eb1 comet length in cultured hippocampal neurons. We co-expressed active GSK-3β^CA^ or GSK-3β^DN^ with EB3-GFP, fixed and stained cells for GFP in 1 DIV neurons (**Fig. 2 E,F**). We did not observe a difference in comet length in neurons expressing GSK-3β^DN^. However, GSK-3β^CA^ caused reductions in comet size (75 % of control).

These experiments suggest that loss of Eb1 from microtubule plus ends could explain the microtubule unbundling that is caused by GSK-3β^CA^ expression, but not GSK-3β^DN^. This indicates that the microtubule disorganisation phenotypes induced by hyper- or in-activation of GSK-3β are caused by different mechanisms.

### GSK-3β hyperactivation leads to microtubule disorganisation via loss of Tau

The microtubule-associated protein Tau is an established GSK-3β phosphorylation target ^45,46^. Upon phosphorylation, Tau detaches from microtubules ^47,48^ and loss of Tau from microtubules leads to decrease of Eb1 at plus ends resulting in microtubule unbundling ^18^. We hypothesised that GSK-3β hyper-activation causes microtubule disorganization by phosphorylating Tau and subsequent Tau detachment from microtubules. To test this hypothesis, we first determined if changes to GSK-3β activity affect Tau association with axonal microtubules. We expressed hyper-active *sggCA* or inactive *sggDN* in a fly line that had GFP-tagged endogenous Tau ^27^, cultured neurons from these flies, and performed live-imaging. In control cells, Tau-GFP exhibited the expected localisation to axonal microtubules (**Fig. 3A**). In contrast, axonal Tau-GFP was substantially reduced in neurons expressing *sggCA*. Expression of *sggDN* did not affect Tau-GFP localisation. We quantified Tau-GFP intensity in the axon and found that *sggCA* expression resulted in a 38% reduction in axons. This shows that GSK3β hyper-activation causes Tau to detach from axonal microtubules and this finding is consistent with the hypothesis that GSK-3β over-activity causes microtubule disorganization through mis-regulation of Tau.

**Figure 3.**
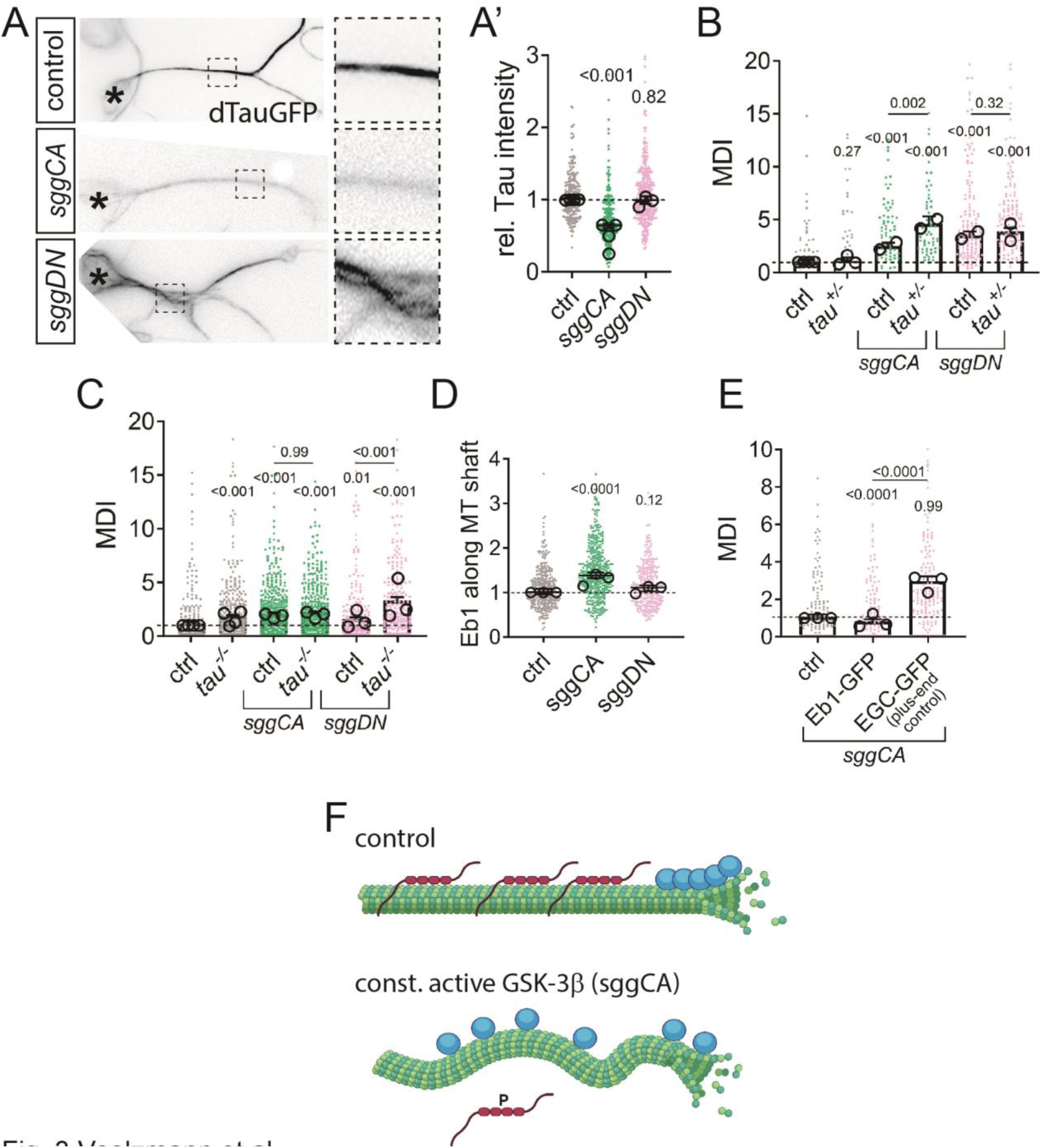
Reduced Eb1 at plus ends through a loss of Tau leads to microtubule unbundling when GSK-3β is hyperactive. **A)** Representative images of primary neurons expressing dTau-GFP (P{Wee}tau[304]) at endogenous levels, alone or in combination with const. active (*UAS-sggCA*) or inactive (*UAS-sggDN*) GSK-3β expression. Asterisks indicate cell bodies, dashed squares are shown as 3.5-fold magnified close-ups beside each image. **A’)** Quantification of relative Tau mean intensity along microtubules for indicated conditions; data are normalised to control. **B, C)** Microtubule curling for primary neurons displaying heterozygous (B) and homozygous (C) *tau^KO^*mutant conditions, alone or in combination with expression of const. active (*UAS-sggCA*) or inactive (*UAS-sggDN*) GSK-3β via elavGal4. **D)** Relative Eb1 intensity along microtubules of neurons at 6 HIV without/with *elav-Gal4*-driven expression of *sggCA* or *sggDN*. **E)** Microtubule curling (MDI) in primary neurons expressing *sggCA* in combination with *UAS-Eb1-GFP* or -*EGC-GFP* (Shot C-terminus). (A’-E) All data were normalised to parallel controls (dashed horizontal lines) and are shown as bar chart with mean ± SEM of at least two independent repeats with 3 technical replicates each; large open circles in graphs indicate mean of independent biological repeats. P-values above data points/bars were obtained with Kruskal-Wallis ANOVA tests. For raw data see S3 Data. **F)** Model view of the results shown here; note that microtubules are depicted (green), blue circles represent Eb1 and red chains Tau; for further explanations see main text and Discussion. Figure generated with Biorender.

To determine if GSK-3β and Tau acted in the same pathway to induce microtubule disorganization, we performed genetic interaction experiments. We used genetically heterozygous flies (*tau^+/-^*), in which one tau allele was mutated to reduce protein levels and measured microtubule disorganisation (**Fig. 3B**). Loss of one *tau* gene copy alone did not cause an increase in microtubule disorganisation. Expression of *sggCA* resulted in increased disorganisation (MDI=2.5), comparable to what we observed above. Expression of *sggCA* in *tau^+/-^* neurons resulted in an exacerbated phenotype (MDI=4.6). Expression of *sggDN* causes increased disorganization (MDI=3.5), but this was not compounded by decreased *tau* expression (MDI=3.9). This suggests that *sggCA,* but not *sggDN* genetically interacts with *tau* and they might act via the same pathway.

To determine if GSK-3β-mediated microtubule disorganisation depended on its interaction with Tau, we expressed constitutively active (*sggCA*) GSK-3β in homozygous *tau* knockout flies (*tau^-/-^*) and measured microtubule disorganisation (**Fig. 3C**). If *sggCA* acts through *tau*, then expression of *sggCA* in *tau^-/-^*neurons will not increase microtubule disorganization compared to *tau^-/-^*neurons not expressing *sggCA*. We performed that experiment and found that loss of *tau^-/-^* neurons did not increase microtubule disorganization in neurons expressing *sggCA* (MDI=1.9 and 2.0 respectively). In contrast, *sggDN* expression in *tau^-/-^*neurons increased microtubule disorganisation compared to *tau^-/-^*neurons from MDI=1.4 to MDI=3.1. Those results further strengthen the model that GSK-3β hyper-activation causes microtubule disorganisation through Tau, and that GSK-3β in-activation induces microtubule disorganisation by a different mechanism.

We previously demonstrated that loss of Tau results in increased Eb1 binding to the microtubule lattice and loss of Eb1 at growing microtubule tips ^18^. Loss of Eb1 at plus ends leads to microtubule curling, because it reduces the ability of the actin-microtubule crosslinker Shot to guide growing microtubules into parallel bundles ^18^. To determine if hyperactive GSK-3β leads to microtubule unbundling by Eb1 relocating from microtubule tips to the shaft, we stained for Eb1 in wt, *sggCA*-, or *sggDN*-expressing neurons, and measured Eb1 at microtubule shafts (**Fig. 3D**). We found that expression of *sggCA* resulted in a 1.3-fold increase of Eb1 at microtubule shafts (areas without comets) compared to control neurons (**Fig. 3D**). Expression of *sggDN* did not change Eb1 levels at microtubule shafts.

Increasing Eb1 levels by over-expressing Eb1-GFP can restore Eb1 to the plus tip and rescue microtubule curling in *tau^-/-^* mutants ^18^. Therefore, we asked if Eb1-GFP overexpression could rescue the microtubule disorganisation phenotype caused by *sggCA* (**Fig. 3E**). Expression of *sggCA* did lead to an increase in microtubule disorganization (MDI=0.8) when Eb1-GFP was co-expressed. This rescue is specific for Eb1, because co-expression of the plus-end binding domain of another protein (Shot, EGC^16^) and *sggCA* still resulted in microtubule disorganisation (MDI=2.92). This shows that GSK-3β hyperactivation causes microtubule disorganization through Tau and Eb1 (**Fig. 3F**).

### Shot and inactive GSK-3β act in the same pathway to cause microtubule unbundling

A key mediator of microtubule bundling is the Spectraplakin Short Stop (Shot) which guides polymerising microtubules into parallel bundles by crosslinking cortical actin and Eb1^16^. Therefore, Shot is a plausible target for GSK-3β misregulation to cause microtubule disorganisation. To determine if GSK-3β and *shot* act in the same pathway, we generated heterozygous *shot^+/-^* flies to reduce *shot* expression, cultured neurons from these animals, and measured microtubule disorganisation (**Fig. 4A**). The MDI of *shot^+/-^* neurons was statistically identical to the MDI in wild type neurons, indicating that even reduced shot levels were sufficient for proper bundling of microtubules. We then expressed *sggCA* or *sggDN* in wild type and *shot^+/-^* neurons and measured microtubule disorganization (**Fig. 4A**). S*ggCA* expression increased microtubule disorganisation significantly more in *shot^+/-^*neurons (MDI=3.88) than in wild type neurons (MDI=2.53). *SggDN* expression also increased microtubule disorganisation more in *shot^+/-^* neurons (MDI=5.73) than in wild type neurons (MDI=2.75). To further test if *sggDN* acts through *shot*, we cultured neurons from *shot* null mutants (*shot^-/-^*) and determined if GSK-3β expression enhanced the microtubule disorganisation beyond the level caused by loss of *shot* (**Fig. 4B**). *Shot* deletion resulted in substantial microtubule disorganisation (MDI=4.14), as expected. However, expression of neither *sggCA* (MDI=3.8) nor *sggDN* (MDI=3.7) increased microtubule disorganisation beyond what was observed in homozygous *shot^-/-^*neurons.

**Figure 4.**
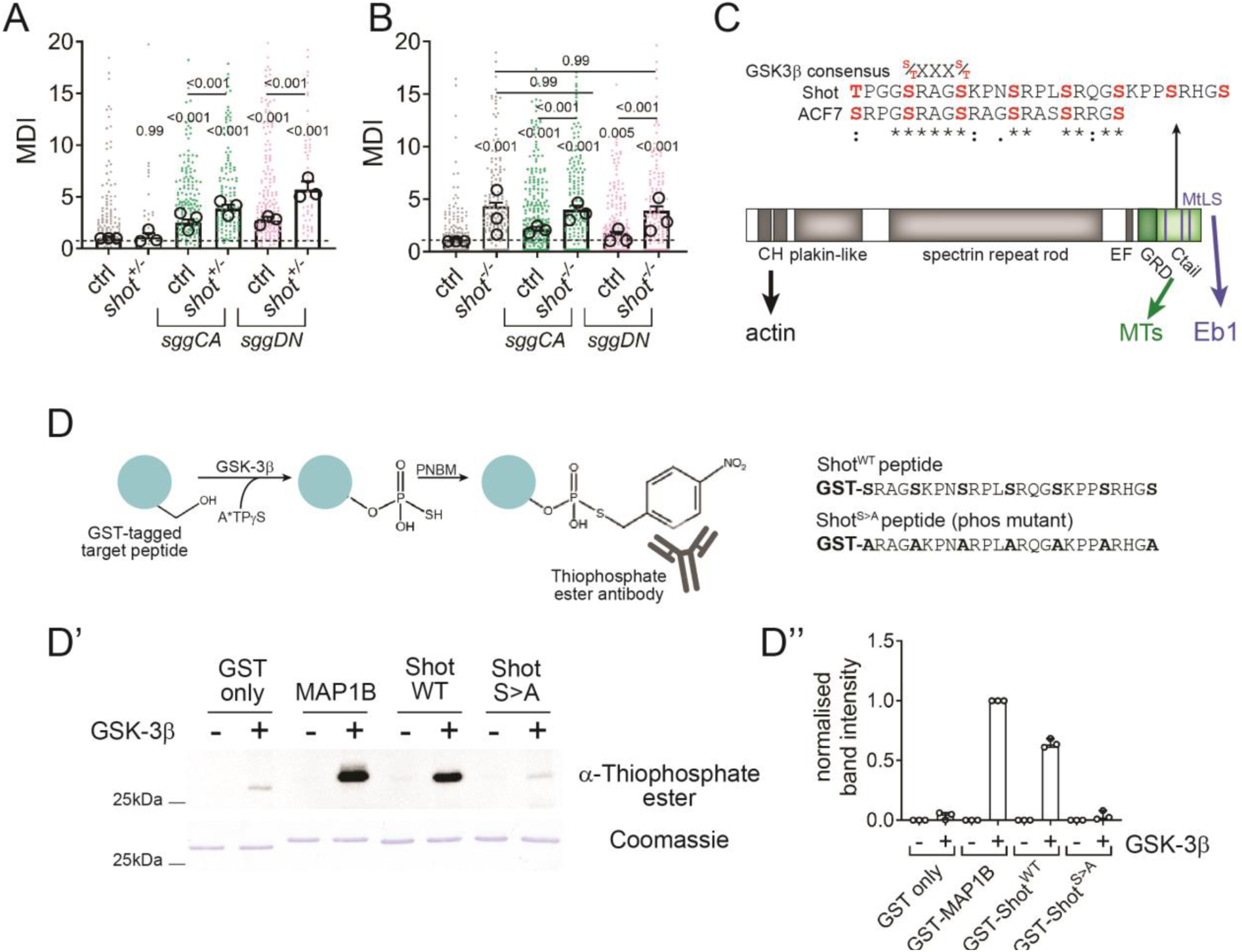
The C-terminus of Shot can be phosphorylated by GSK-3β. **A, B)** Microtubule curling for primary neurons displaying heterozygous (A) and homozygous (B) *shot*^3^ mutant conditions, alone or in combination with expression of const. active (*UAS-sggCA*). **C)** Schematic representation of Shot; highlighting the verified ACF7 and putative Shot GSK-3β target site (consensus S/T S/T) in the C-terminal microtubule binding region. **D)** Workflow of the in vitro thiophosphorylation kinase assay and Shot^WT^ and phosphodeficient Shot^S>A^ peptide sequence. D’) Representative blot where phosphorylated peptides are labelled with α-Thiophosphate ester of GST control, positive control (GSK-3β target site of MAP1B, ERLSPAKSPSLSPSPPSPIEKT^51^), Shot^WT^ and Shot^S>A^ with or without GSK-3β. D’’) Quantification of mean band intensity. See Fig. 4-S1 for full blots. For raw data see S4 Data.

The observations that *shot* knockout alone achieved maximal disorganisation and that disorganisation was not enhanced by concurrent expression of *sggCA* or *sggDN*, suggest that GSK-3β manipulations that increase bundling are mechanistically linked to *shot* activity. These results are also strong evidence that Shot and GSK-3β act in the same pathway and support the hypothesis that Shot is the relevant GSK-3β substrate.

### GSK-3β phosphorylates Shot in its Eb1/microtubule-binding region

GSK-3β phosphorylates serine or threonine residues of defined consensus motifs ^49^. In the mammalian Shot homologue ACF7, GSK-3β phosphorylates a cluster of serines that represent a canonical GSK-3β consensus motif (**Fig. 4C**)^50^. They are located in between the C-terminal microtubule binding domain (Gas domain) and Eb1 protein-binding SxIP sites. We identified a comparable cluster of putative GSK-3β phosphorylation sites in the C-terminus of Shot by sequence analysis (residues 5040 – 5064 of Shot-PE, **Figs. 4C, 4-S1A**).

To determine if GSK-3β phosphorylates these residues, we generated a GST-tagged peptide of the wild type cluster (Shot^WT^) and a corresponding peptide with every serine residue mutated to alanine (Shot^S>A^). We separately generated a peptide of a human MAP1B sequence that is phosphorylated by GSK-3β (ERLSPAKSPSLSPSPPSPIEKT; ^51^) to use as a positive control. We tested these peptides in an in vitro GSK-3β thiophosphorylation assay ^52^ (**Fig. 4D**).

Phosphorylation of GST was minimal, while the MAP1B peptide exhibited strong phosphorylation. Shot^WT^ was substantially phosphorylated, identifying this fragment of Shot as a GSK-3β phosphorylation target. In contrast, Shot^S>A^ was not phosphorylated, showing that phosphorylation of the Shot sequence required the cluster of serine residues (**Figs. 4D’,D’’, 4-S1B**). These data show that GSK-3β can phosphorylate the C-terminal tail of Shot.

### Shot phosphorylation is required to maintain axonal microtubule organisation

Because GSK-3β can phosphorylate Shot, we asked if Shot phosphorylation was important for axonal microtubule organisation. We modified the *shot* locus and generated three different fly lines where the final three exons were replaced with a merged single exon that also encoded a C-terminal GFP (**Figs. 5A, 5-S1**): i) a wild type line, ii) a phosphomimetic line where the seven serine residues in the GSK-3β phosphorylation motif were replaced with aspartic acids (**Fig. 4C**), and iii) a phosphonull line where the same serine residues were replaced by alanine. Neither the phosphomimetic, nor the phosphonull Shot can be tuned by GSK-3β phosphorylation in this C-terminal serine-rich cluster.

**Figure 5.**
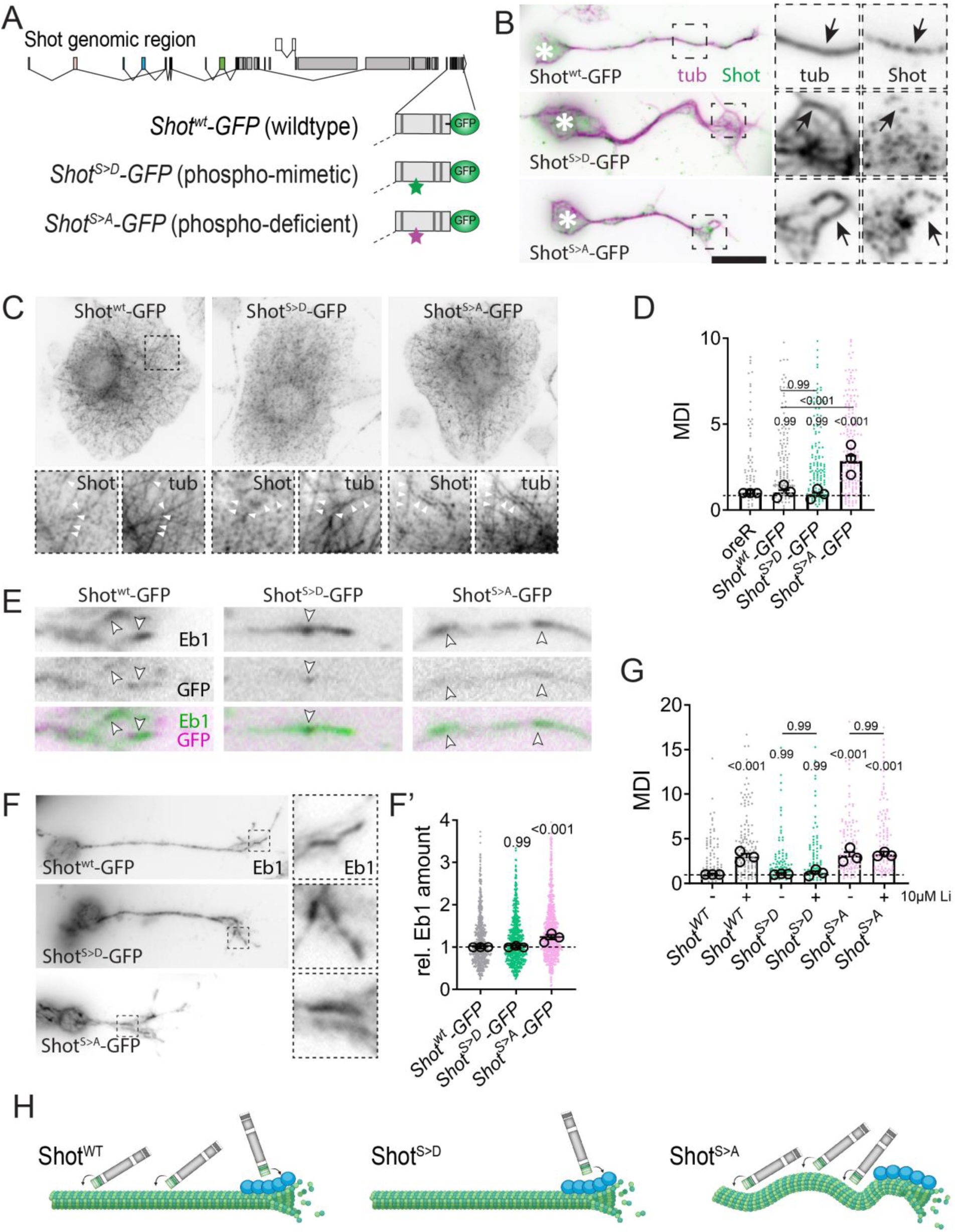
Phosphodeficient but not -mimetic Shot leads to microtubule unbundling. **A)** Schematic representation of genomic engineering approach replacing C-terminal exons of Shot to generate endogenous GFP-tagged Shot CRISPR lines, as either wildtype (*Shot^WT^*), phosphomimetic (*Shot^S>D^*) or -deficient (*Shot^S>A^*) variants. **B, C, E, F)** Representative images of *Drosophila* primary neurons (B, E, F) and non-neuronal, fibroblast-like cells (C) cultured for 6HIV of *Shot^WT^*, *Shot^S>D^* or *Shot^S>A^* labelled for tubulin (purple) and Shot (green, anti-GFP) in (B, C), Eb1 (green) and GFP (magenta) in (E) and Eb1 in (F). Asterisks indicate cell bodies, dashed squares in are shown as 3.5-fold magnified close-ups beside (B, F) or underneath (C) each image with tubulin and GFP (Shot) channels, arrows/white arrowheads point at microtubules in B,C and at Eb1 comets in E, F. **F’)** Quantification of relative Eb1 amounts at plus ends for conditions indicated. **D, G)** Quantification of microtubule unbundling (MDI) obtained from embryonic primary neurons with conditions indicated below. **G)** Primary neurons were treated for the entire 6HIV with 10 mM Li or vehicle. **H)** Cartoons, generated by Biorender, illustrating current hypothesis how phosphorylation by GSK-3β affects Shot’s ability to bind plus ends or microtubules. For raw data see S5 Data.

We cultured primary neurons from homozygous embryos of each line for 6 HIV and stained for microtubules and GFP to enhance the Shot-GFP signal (**Fig. 5B**). In all conditions, Shot-GFP was expressed at low levels and exhibited a punctate distribution in axons. Shot^WT^-GFP puncta frequently colocalized with microtubules, although the low intensity of Shot-GFP puncta prevented systematic colocalization quantification. In contrast, phosphomimetic Shot^S>D^-GFP exhibited little colocalization with microtubules. Phosphodeficient Shot^S>A^-GFP localised to microtubules, comparable to Shot^wt^-GFP.

We performed complementary experiments in *Drosophila* fibroblast-like cells co-cultured with the primary neuron population and found comparable results. Shot^WT^-GFP and Shot^S>A^-GFP colocalized with microtubules, but Shot^S>D^ did not (**Fig. 5C**). These results support the hypothesis that Shot’s ability to bind microtubules is regulated by GSK-3β-mediated Shot phosphorylation.

We next determined if the phosphorylation state of Shot affects microtubule bundling. We cultured neurons from the same lines for 6 HIV, stained for microtubules, and measured microtubule disorganisation by MDI (**Fig. 5D**). Neurons from homozygous *shot^WT^* and *shot^S>D^* embryos exhibited no substantial microtubule disorganisation. In contrast, neurons cultured from phosphodeficient *shot^S>A^* mutants showed significant microtubule disorganisation (MDI=2.9).

Even though Shot^S>A^-GFP, in contrast to Shot^S>D^-GFP, binds microtubule efficiently, the phosphodeficient S>A mutation leads to microtubule disorganisation. This shows that Shot binding of the microtubule shaft is not linked with microtubule disorganisation. An explanation for microtubule unbundling in phospho-deficient Shot^S>A^-GFP neurons could therefore come from Shot’s interaction with Eb1. If Shot cannot bind Eb1, it loses its ability to guide polymerising microtubules into parallel bundles ^16^. To test if shot^S>A^-GFP leads to microtubule unbundling due to reduced Shot–Eb1 interaction, we cultured neurons from the three Shot-GFP lines for 6 HIV and stained for Eb1. We found that Shot^S>D^-GFP, but not Shot^S>A^-GFP, co-localised with Eb1, (**Fig. 5E**; low intensity of shot-GFP puncta prevented systematic colocalization quantification). These results support the hypothesis that Shot^S>A^-GFP cannot efficiently bind Eb1.

Previous work found that loss of Shot–Eb1 interaction causes increased Eb1 comet sizes because microtubule-bound Shot slows microtubule polymerisation ^16^. To determine if the phosphorylation state of Shot affected Eb1 binding to microtubule plus ends, we quantified the comet size in each condition. Shot^S>D^-GFP expressing neurons did not show any effect on comet size (**Fig. 5F,F’**). In contrast, Eb1 comets were significantly larger in the presence of phosphodeficient Shot^S>A^-GFP. Taken together, these results support the hypothesis that Shot– Eb1 interaction is lost when Shot is dephosphorylated, which leads to a loss of Shot-mediated microtubule guidance and microtubule disorganisation.

To determine if Shot and GSK-3β are two components of the same mechanism or independent pathways, we use the genetically modified *shot* fly lines. If the effects of GSK-3β inhibition and Shot manipulation on microtubule unbundling are additive, it would suggest they operate through separate pathways. However, if GSK-3β inhibition does not increase microtubule disorganisation in the *shot* phospho-mutants, it indicates they are acting via same mechanisms. We cultured primary neurons from Shot^WT^-GFP, Shot^S>A^-GFP and Shot^S>D^-GFP animals for 6 HIV in the presence or absence of the GSK-3β inhibitor lithium (10 mM LiCl). Incubation of Shot^WT^ neurons with 10 mM LiCl lead to a three-fold increase in microtubule unbundling (**Fig. 5G**). In contrast, LiCl treatment did not increase microtubule unbundling in Shot^S>A^-GFP (MDI=3.14 vs 3.23) and, importantly, did not cause microtubule disorganisation in Shot^S>D^-GFP (MDI=1.04 vs 1.2). Therefore, unbundling of microtubules by GSK-3β inhibition and loss of Shot phosphoregulation are not additive. These observations are strong evidence that the microtubule unbundling caused by GSK-3β inhibition is mediated by Shot.

## Discussion

### A novel role for GSK-3β in maintaining axonal microtubule bundles

We describe a new, evolutionary conserved mechanistic model of how GSK-3β maintains parallel microtubule bundles in neurons. We found that GSK-3β activity must be tightly balanced to maintain parallel bundles of microtubules; hyper- or inactivation of GSK-3β leads to curling of microtubules in fly and rat neurons. GSK-3β regulates microtubule bundling through the microtubule binding proteins Tau and Shot **(Fig. 6)**, two microtubule regulators that maintain microtubule bundling through Shot/Eb1 mediated guidance of microtubule polymerisation ^18^. Shot guides polymerising microtubules into parallel bundles by crosslinking actin and Eb1 ^16^. The Shot/Eb1 guidance mechanism breaks in different ways when GSK-3β is inhibited or hyperactive. In both situations, Shot-mediated guidance of polymerising microtubules is reduced, leading to aberrant growth of unbundled microtubules towards the plasma membrane ^18^. GSK-3β hyperactivity causes Tau to detach from microtubules, which results in Eb1 binding to microtubule shafts. Sequestration of Eb1 to microtubule shafts reduces Eb1 that is available to guide polymerising microtubules into properly formed bundles by reducing Shot/Eb1 interaction. GSK-3β inhibition reduces microtubule guidance by Shot by reducing the affinity of Shot for Eb1.

**Figure 6.**
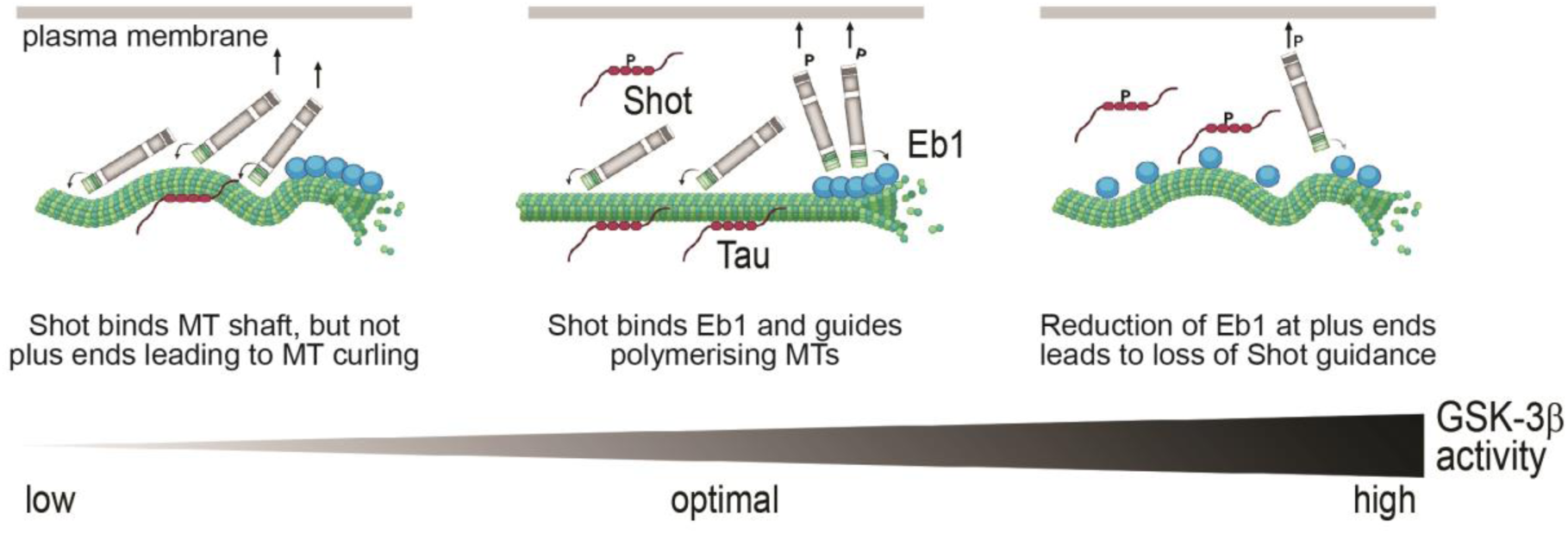
GSK-3β activity needs to be tightly regulated to control microtubule bundling - a mechanistic model consistent with reported data. Moderate GSK-3β activity levels are optimal to maintain microtubule bundles: Unphosphorylated Tau binds the microtubule shaft and prevents Eb1 from binding the lattice. Low levels of Shot phosphorylation ensure that Shot can interact with Eb1 and guide polymerising microtubules (MTs) into parallel bundles. Low activity of GSK-3β leads to a strong association of Shot with the microtubule shaft and reduction of Eb1 binding, thereby losing its ability to guide polymerising microtubules. Consequently, microtubules are disorganised. Hyper-activity of GSK-3β causes a loss of Tau and increase of Eb1 along the microtubule shaft. Eb1 comet size is reduced, and Shot is not able to efficiently bind plus ends and guide polymerising microtubules.

### GSK-3β activity must be tightly balanced to maintain microtubule bundles in health axons

Microtubules provide essential structural support for axons and are the substrate for all long-range vesicle transport. Unsurprisingly, proper maintenance of microtubule bundles is required for neuronal development and health. Defects in microtubule bundles can trigger axonal decay, which causes axon swellings that impair vesicle transport and action potential propagation, key features of axonopathies ^4^. Pathological axon swellings in which microtubules bundles have disintegrated into loops or waves have been observed in ageing, after injury, and in animal models of axonopathies ^3^. GSK-3β is the key coordinator of the parallel microtubule architecture in axons. Therefore, unbundling of axonal microtubules is a likely mechanism by which impaired GSK-3β activity leads to neuronal decay.

The present study shows that GSK-3β activity needs to be tightly balanced to maintain bundled axonal microtubules. GSK-3β hyperactivity impairs neuronal development and regeneration and drives key features of Alzheimer’s Disease (Aβ production and accumulation, formation of toxic tau species, pro-inflammatory and LTP impairments; reviewed by Lauretti et al. ^53^). A recent study found that genetic variation resulting in GSK-3β loss-of-function leads to neurodevelopmental disorders, developmental delay and individuals exhibiting core features of autism ^54^. Inhibition of GSK-3β or treatment with GSK-3β inhibitors can lead to limited regenerative outgrowth ^55^, reduced outgrowth ^56^, and defects in neuronal plasticity, such as neuromuscular junction defects in flies ^57^. GSK-3β is considered a highly promising therapeutic target ^58^. However, global GSK-3β inhibitors have failed due to lack of benefit or toxicity. GSK-3β is a required kinase, therefore excessive inhibition of GSK-3β could negatively impact its normal functions, such as the coordination of microtubule integrity which we observe here. The proper balance between GSK-3β activation and inhibition is required to drive normal development where multiple complex regulatory layers ensure that GSK-3β regulates its substrates at the appropriate time and location. However, we do not yet fully understand GSK-3β’s spatiotemporal activity patterns yet. Furthermore, the regulation of microtubule networks by GSK-3β is complex and the impact of the kinase is highly dependent on individual targets, e.g. active GSK-3β is reported to promote microtubule dynamics by phosphorylating MAP1b ^51^, inhibition of GSK-3β increases microtubule stability via Tau, CRMP2 or APC (reviewed in ^7^. For others GSK-3β targets such as Clasp ^59^ or Shot (this study) GSK-3β activity levels determine where they bind microtubules, at the microtubule shaft or plus end. A full understanding of this complex regulatory network will require highly tractable genetic approaches with novel tools that elucidates specific activity patterns of GSK-3β. This is the crucial next step to explore how the complexity of GSK-3β signalling is coordinated in time and space to achieve precise control over a wide range of cellular processes during neural development.

Our data support a model where phosphorylation by GSK-3β balances two of Shot’s key functions – binding/stabilising microtubules directly and guiding microtubule polymerisation by binding Eb1 ^16^. Wu et al. ^50^ found that microtubule binding of the Spectraplakin ACF7 is regulated by GSK-3β, where ACF7 detaches from microtubules upon phosphorylation. We similarly observed that *Drosophila* Shot–microtubule association was reduced with phosphomimetic Shot. In addition, we found that Shot’s interaction with Eb1 depends on the phosphorylation state of Shot, where phosphomimetic Shot can still bind Eb1 despite being detached from microtubules. This switch could be the mechanism by which signalling coordinates and controls two key functions of Shot: 1) stabilising and protecting microtubules when it is unphosphorylated and 2) guiding polymerising microtubules into parallel bundles. This is reminiscent of a similar mechanism that was described for Clasp which contains two GSK-3β motives ^59,60^. Here, the number of phosphorylated residues determine whether it associates with the plus end, the microtubule shaft or shows no interaction with microtubules at all. In contrast to Shot, complete phosphorylation of all GSK3 sites disrupts both the plus end-tracking and the lattice-binding activities of CLASP2 ^59^. While Wu et al. did not directly analyse ACF7’s interaction with Eb proteins, they observed that directionality of microtubule growth was largely random in ACF7 KO cells. Expression of wildtype ACF7, but not phosphodeficient ACF7(S:A) rescued properly directed microtubule growth. This suggests that the role of GSK-3β in the spectraplakin/Eb interaction and guidance of microtubule polymerisation is evolutionary conserved. It is not yet clear how phosphorylation mediates a shift between the microtubule- and Eb1-bound states of Shot. Our structural in silico analysis using AlphaFold 2 suggests that a key GSK-3β target cluster is in a linker region between Shot’s three SxIP sites that might sit in between two associated Eb1 molecules (**Fig.5-S2**) Increasing the negative charge in this linker region may lead to a loss of microtubule affinity and strengthening interactions with Eb1.

### Microtubule organisation in neurodegenerative diseases and ageing

Our findings are important for neurodegenerative diseases in which GSK-3β is hyperactive and leads to hyperphosphorylation of Tau and the formation of neurofibrillary tangles, such as Alzheimer’s and Parkinson’s Disease. Much emphasis has been placed on those aggregates. However, work in flies and mouse models suggested that loss of Tau alone does not cause severe phenotypes ^23^ ^61^ apart from mild behavioural and motor deficits (e.g. ^62,63^. Therefore, loss of endogenous Tau function may have another important effect. We previously showed that Tau plays an important role in microtubule polymerisation and bundle organisation by keeping Eb1 off the microtubule shaft ^18^. This is consistent with the observation in this study that GSK-3β-mediated detachment of Tau from microtubules leads to microtubule unbundling. However, we find that when GSK-3β is hyperactive Shot loses binding microtubules and Tau. Shot and Tau are functionally redundant in the context of stabilising microtubules where the combined loss of both microtubule binding proteins exceeds the impact of loss of just one exponentially, leading to severe microtubule instability, impaired transport and synapse defects ^64^. Therefore, the simultaneous loss of both, Tau and Shot, from microtubules could potentially have a detrimental impact on neuronal cell biology leading to decay, in addition to pathological microtubule unbundling. We therefore propose the combined loss of Tau and Shot as new mode how hyperactive GSK-3β leads to neuronal decay in neurodegenerative diseases.

The loss of microtubule organization caused by GSK-3β dysregulation might be not only be relevant in neurodegenerative diseases but in aging, where many neuronal axons are lost ^65^. A recent study found that Tau, Shot, and EB1 are essential for maintaining microtubules in ageing axons. Loss of these proteins in ageing neurons leads to the decay of microtubule bundles, which subsequently cause a decline in axons and synaptic terminals ^66^. Notably, GSK-3β expression increases with age ^67,68^. Therefore, mis-regulation of GSK-3β activity could be an underlying cause for the loss of microtubule regulators and therefore microtubule integrity as we get older.

GSK-3β is a highly promising therapeutic target for various neurological disorders, from Alzheimer’s, Parkinson’s and Huntington’s disease to neurodevelopmental and mood disorder. However, trials involving global GSK-3β inhibitors have largely failed. We propose a model that explains why global inhibition of GSK-3β has not been successful. Deviating even slightly from optimal levels of GSK-3β leads to problems with neuronal function and morphology. Thus, effective interventions will require precise spatial and temporal regulation of GSK-3β activity within nerve cells.

## Supporting information

raw data

## Acknowledgements

This work was made possible through support by the Leverhulme Trust (ECF-2017-247), Academy of Medical Sciences Springboard Award (SBF008\1140) and Royal Society Research Grant (RGS\R2\222151) to I.H., AV was funded by startup funding by University of Hull and the BBSRC to Andreas Prokop (BB/I002448/1, BB/P020151/1, BB/L000717/1, BB/M007553/1). This work was supported by National Institutes of Health (NIH) award R35GM153259 to M.B. The Manchester Bioimaging Facility microscopes used in this study were purchased with grants from the BBSRC, The Wellcome Trust and The University of Manchester Strategic Fund. Work on this project benefited from the Manchester Fly Facility, established through funds from the University and the Wellcome Trust (087742/Z/08/Z). We would like to thank Peter O’Toole, Grant Calder and Karen Hogg at the York Bioscience Technology Facility for assistance with microscopy. We thank Andreas Prokop for his support and helpful discussions throughout this project, Hiro Ohkura for kindly providing DmEb1 antibody. Stocks obtained from the Bloomington *Drosophila* Stock Center (NIH P40OD018537) were used in this study.

---

**Figure 4-S1.**
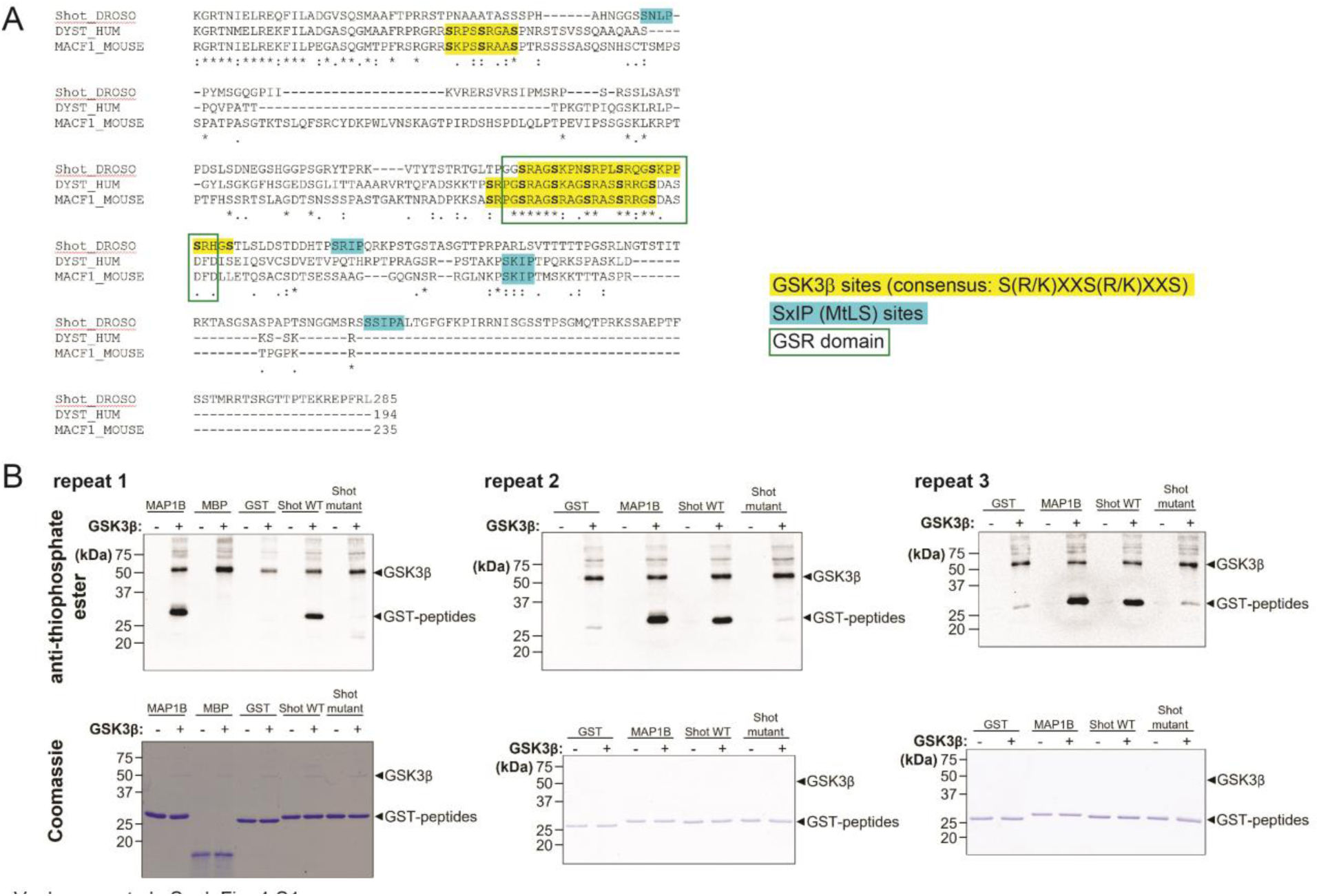
Phosphorylation of the C-terminus of Shot by GSK-3β. **A)** Sequence alignment of the C-termini of *Drosophila* Shot (shot-RE FBpp0086744), human Dystonin (DST-001, ENST00000370788) and mouse MACF1 (Macf1-201, ENSMUST00000084301) highlighting C-terminal domains (square), EB-binding SxIP sites (blue) and GSK-3β consensus motifs (yellow). **B)** Western blots of Coomassie gels of three repeats of the in vitro thiophosphorylation kinase assay Schematic representation of Shot; highlighting the verified ACF7 and putative Shot GSK-3β target site (consensus S/T S/T) in the C-terminal microtubule binding region. GST only (negative control), GSK-3β target site of MAP1B (positive control; ERLSPAKSPSLSPSPPSPIEKT ^51^), Shot^WT^ and Shot^S>A^ (see Fig. 4D) with or without GSK-3β. Coomassie shows protein loading, blots show phosphorylated peptides are labelled with α-Thiophosphate ester. For raw data see S4 Data.

**Figure 5-S1.**
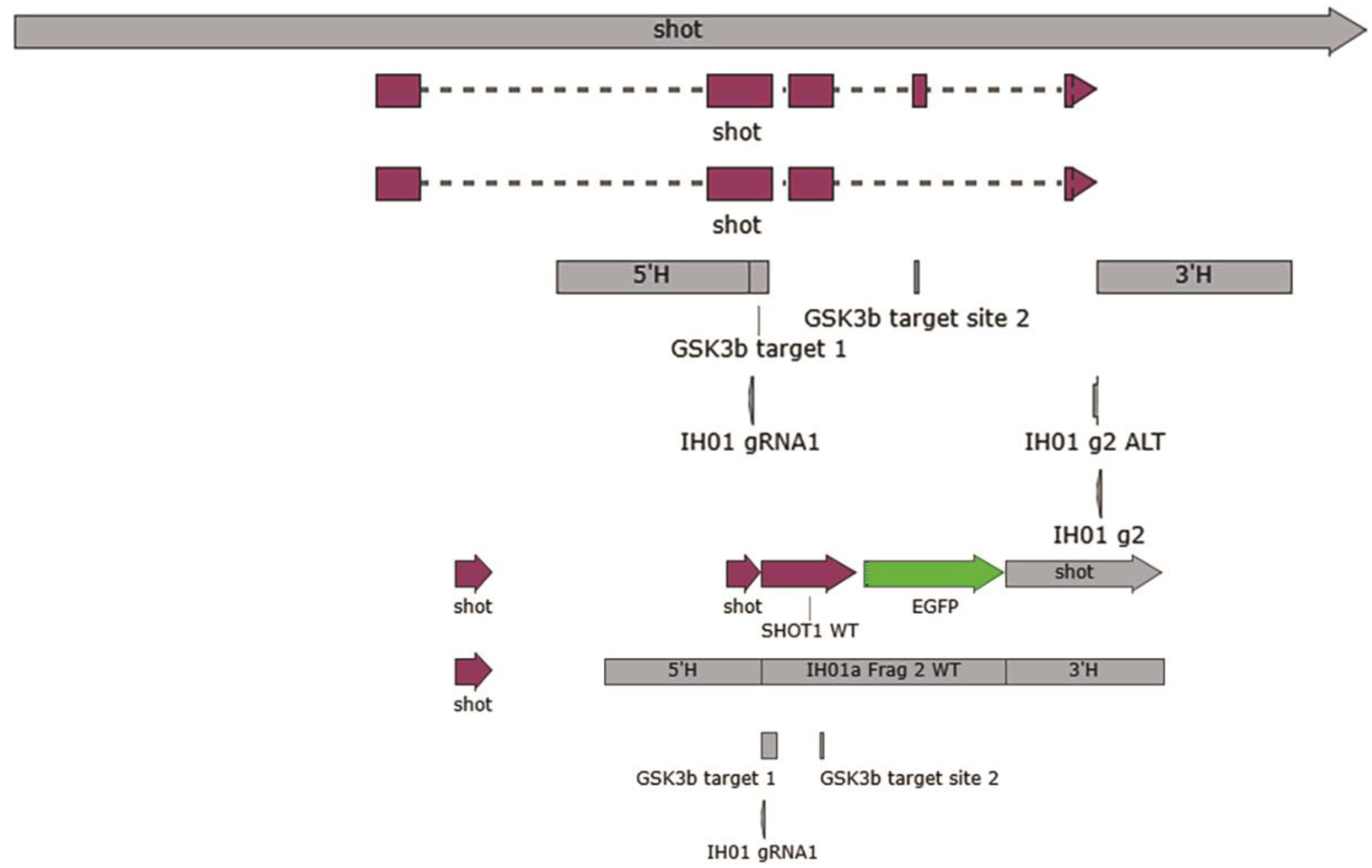
CRISPR/Cas9 mediated genomic targeting strategy to generate wildtype, phospho-mutant and -mimic Shot alleles. The five C-terminal exons of Shot, 5’ and 3’ homology arms and targeting sites for guide RNAs are depicted as well as an exemplar homologous repair construct (Shot^WT^).

**Figure 5-S2.**
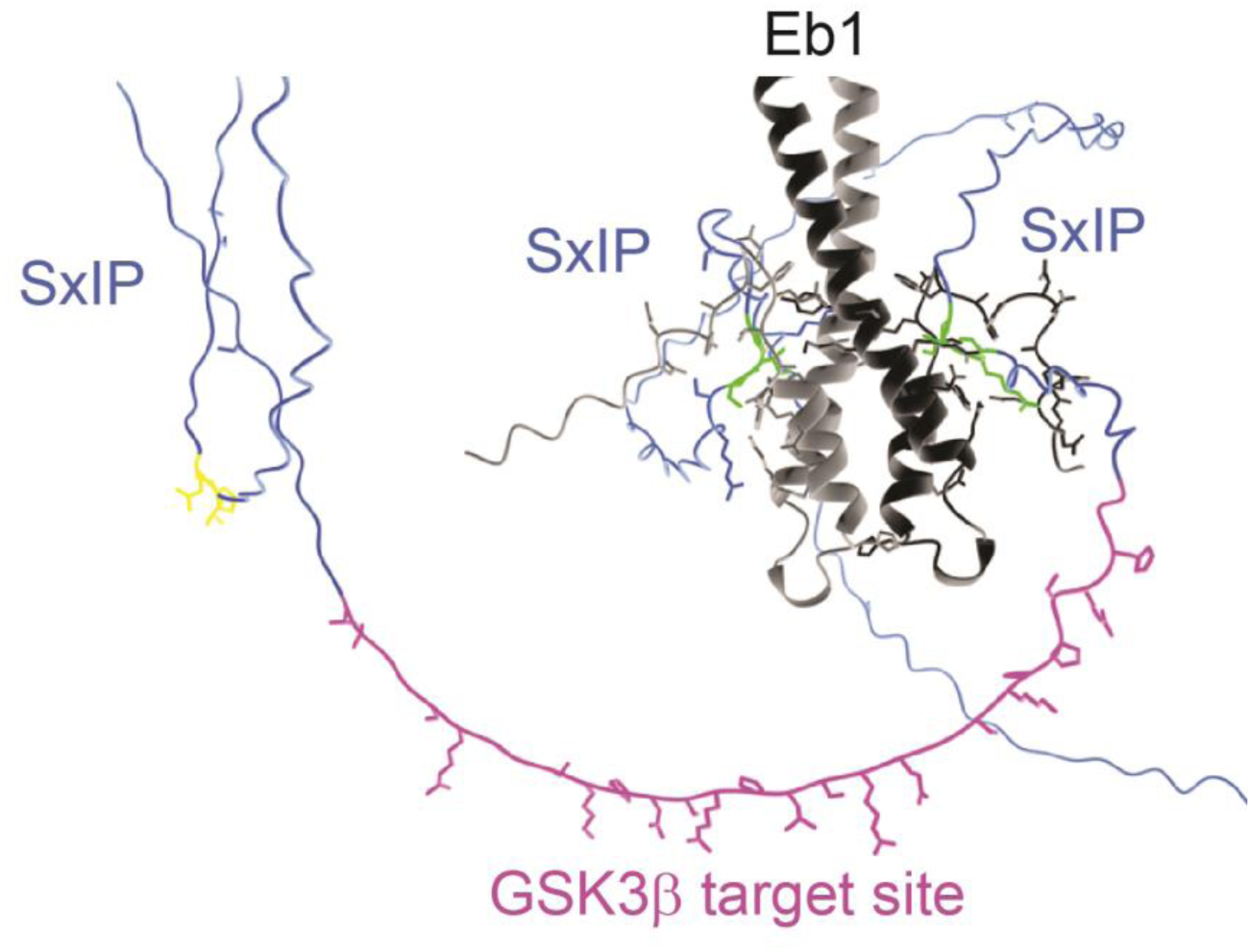
AlphaFold 2 model of Shot C-terminus and Eb1. The GSK-3β target sites is located in between the three Eb1-binding SxIP sites.

## Notes

### Competing Interest Statement

The authors have declared no competing interest.

### Summary of Updates

We edited the text throughout to make the manuscript more concise. We updated all Figures and added a summary Figure 6 and supplementary Figures 4-S1, 5-S1 and 5-S2. Affiliations were updated as well.

## References

1 Kapitein, L. C. & Hoogenraad, C. C. Building the neuronal microtubule cytoskeleton. Neuron 87, 492–506 (2015). 10.1016/j.neuron.2015.05.046 S0896-6273(15)00513-9 [pii]

2 Bentley, M. & Banker, G. The cellular mechanisms that maintain neuronal polarity. Nat Rev Neurosci 17, 611–622 (2016). 10.1038/nrn.2016.100

3 Hahn, I., Voelzmann, A., Liew, Y.-T., Costa-Gomes, B. & Prokop, A. The model of local axon homeostasis - explaining the role and regulation of microtubule bundles in axon maintenance and pathology Neural Dev 14, 10.1186/s13064-13019-10134-13060 (2019).

4 Prokop, A. A common theme for axonopathies? The dependency cycle of local axon homeostasis. Cytoskeleton (Hoboken) 78, 52–63 (2021). 10.1002/cm.21657

5 Voelzmann, A., Hahn, I., Pearce, S., Sánchez-Soriano, N. P. & Prokop, A. A conceptual view at microtubule plus end dynamics in neuronal axons. Brain Res Bulletin 126, 226–237 (2016). 10.1016/j.brainresbull.2016.08.006

6 Luo, J. The role of GSK3beta in the development of the central nervous system. Front Biol (Beijing) 7, 212–220 (2012). 10.1007/s11515-012-1222-2

7 Hur, E. M. & Zhou, F. Q. GSK3 signalling in neural development. Nat Rev Neurosci 11, 539–551 (2010). 10.1038/nrn2870

8 Llorens-Martin, M., et al. GSK-3beta overexpression causes reversible alterations on postsynaptic densities and dendritic morphology of hippocampal granule neurons in vivo. Mol Psychiatry 18, 451–460 (2013). 10.1038/mp.2013.4

9 Xing, B., Li, Y. C. & Gao, W. J. GSK3beta Hyperactivity during an Early Critical Period Impairs Prefrontal Synaptic Plasticity and Induces Lasting Deficits in Spine Morphology and Working Memory. Neuropsychopharmacology 41, 3003–3015 (2016). 10.1038/npp.2016.110

10 Rizk, M., et al. Deciphering the roles of glycogen synthase kinase 3 (GSK3) in the treatment of autism spectrum disorder and related syndromes. Mol Biol Rep 48, 2669–2686 (2021). 10.1007/s11033-021-06237-9

11 Jorge-Torres, O. C., et al. Inhibition of Gsk3b Reduces Nfkb1 Signaling and Rescues Synaptic Activity to Improve the Rett Syndrome Phenotype in Mecp2-Knockout Mice. Cell Rep 23, 1665–1677 (2018). 10.1016/j.celrep.2018.04.010

12 Duda, P., et al. Targeting GSK3 signaling as a potential therapy of neurodegenerative diseases and aging. Expert Opin Ther Targets 22, 833–848 (2018). 10.1080/14728222.2018.1526925

13 Chokhawala, K., Lee, S. & Saadabadi, A. in StatPearls (2025).

14 Arciniegas Ruiz, S. M. & Eldar-Finkelman, H. Glycogen Synthase Kinase-3 Inhibitors: Preclinical and Clinical Focus on CNS-A Decade Onward. Front Mol Neurosci 14, 792364 (2021). 10.3389/fnmol.2021.792364

15 Bottenberg, W., et al. Context-specific requirements of functional domains of the Spectraplakin Short stop in vivo. Mech Dev 126, 489–502 (2009).

16 Alves-Silva, J., et al. Spectraplakins promote microtubule-mediated axonal growth by functioning as structural microtubule-associated proteins and EB1-dependent +TIPs (Tip Interacting Proteins). J. Neurosci 32, 9143–9158 (2012). 10.1523/JNEUROSCI.0416-12.2012 32/27/9143 [pii]

17 Qu, Y., Hahn, I., Webb, S. E. D., Pearce, S. P. & Prokop, A. Periodic actin structures in neuronal axons are required to maintain microtubules. Mol Biol Cell 28 296–308 (2017). 10.1091/mbc.e16-10-0727

18 Hahn, I., et al. Tau, XMAP215/Msps and Eb1 co-operate interdependently to regulate microtubule polymerisation and bundle formation in axons. PLoS Genet 17, e1009647 (2021). 10.1371/journal.pgen.1009647

19 Liew, Y.-T., Voelzmann, A., Shields, S., Sánchez-Soriano, N. & Prokop, A. Understanding roles of microtubules during kinesin-1-linked axon degeneration. (in preparation)

20 Hazelett, D. J., Bourouis, M., Walldorf, U. & Treisman, J. E. decapentaplegic and wingless are regulated by eyes absent and eyegone and interact to direct the pattern of retinal differentiation in the eye disc. Development 125, 3741–3751 (1998). 10.1242/dev.125.18.3741

21 Bourouis, B. Targeted increase in Shaggy activity levels blocks wingless signaling. Genesis 34, 99–102 (2002).

22 Heitzler, P. & Simpson, P. The choice of cell fate in the epidermis of Drosophila. Cell 64, 1083–1092 (1991). 10.1016/0092-8674(91)90263-x

23 Burnouf, S., et al. Deletion of endogenous Tau proteins is not detrimental in Drosophila. Sci Rep 6, 23102 (2016). 10.1038/srep23102 srep23102 [pii]

24 Kolodziej, P. A., Jan, L. Y. & Jan, Y. N. Mutations that affect the length, fasciculation, or ventral orientation of specific sensory axons in the Drosophila embryo. Neuron 15, 273–286 (1995).

25 Sánchez-Soriano, N., et al. Mouse ACF7 and Drosophila Short stop modulate filopodia formation and microtubule organisation during neuronal growth. J Cell Sci 122, 2534–2542 (2009). 10.1242/jcs.046268 122/14/2534 [pii]

26 Luo, L., Liao, Y. J., Jan, L. Y. & Jan, Y. N. Distinct morphogenetic functions of similar small GTPases: Drosophila Drac1 is involved in axonal outgrowth and myoblast fusion. Genes Dev. 8, 1787–1802 (1994).

27 Clyne, P. J., Brotman, J. S., Sweeney, S. T. & Davis, G. Green fluorescent protein tagging Drosophila proteins at their native genomic loci with small P elements. Genetics 165, 1433–1441 (2003).

28 Stone, M. C., Roegiers, F. & Rolls, M. M. Microtubules have opposite orientation in axons and dendrites of Drosophila neurons. Mol Biol Cell 19, 4122–4129 (2008). 10.1091/mbc.E07-10-1079 E07-10-1079 [pii]

29 Osterwalder, T., Yoon, K. S., White, B. H. & Keshishian, H. A conditional tissue-specific transgene expression system using inducible GAL4. Proc Natl Acad Sci U S A 98, 12596–12601 (2001).

30 Nicholson, L., et al. Spatial and temporal control of gene expression in Drosophila using the inducible GeneSwitch GAL4 system. I. Screen for larval nervous system drivers. Genetics 178, 215–234 (2008). 10.1534/genetics.107.081968

31 Gratz, S. J., et al. Highly specific and efficient CRISPR/Cas9-catalyzed homology-directed repair in Drosophila. Genetics 10.1534/genetics.113.160713 (2014). 10.1534/genetics.113.160713

32 Port, F. & Bullock, S. L. Augmenting CRISPR applications in Drosophila with tRNA-flanked sgRNAs. Nat Methods 13, 852–854 (2016). 10.1038/nmeth.3972

33 Prokop, A., Küppers-Munther, B. & Sánchez-Soriano, N. Using primary neuron cultures of Drosophila to analyse neuronal circuit formation and function. The making and un-making of neuronal circuits in Drosophila 69, 225–247 (2012). 10.1007/978-1-61779-830-6_10

34 Kaech, S. & Banker, G. Culturing hippocampal neurons. Nat Protoc 1, 2406–2415 (2006). 10.1038/nprot.2006.356

35 Rogers, S. L., Rogers, G. C., Sharp, D. J. & Vale, R. D. Drosophila EB1 is important for proper assembly, dynamics, and positioning of the mitotic spindle. J Cell Biol 158, 873–884 (2002).

36 Elliott, S. L., Cullen, C. F., Wrobel, N., Kernan, M. J. & Ohkura, H. EB1 is essential during Drosophila development and plays a crucial role in the integrity of chordotonal mechanosensory organs. Mol Biol Cell 16, 891–901 (2005).

37 Sánchez-Soriano, N., et al. Drosophila growth cones: a genetically tractable platform for the analysis of axonal growth dynamics. Dev Neurobiol 70, 58–71 (2010).

38 Qu, Y., et al. Efa6 protects axons and regulates their growth and branching by inhibiting microtubule polymerisation at the cortex. eLife 8, e50319 (2019).

39 Qu, Y., et al. Re-evaluating the actin-dependence of spectraplakin functions during axon growth and maintenance. Dev Neurobiol 82, 288–307 (2022). 10.1002/dneu.22873

40 Bourouis, M., et al. An early embryonic product of the gene shaggy encodes a serine/threonine protein kinase related to the CDC28/cdc2+ subfamily. EMBO J 9, 2877–2884 (1990). 10.1002/j.1460-2075.1990.tb07477.x

41 Stambolic, V., Ruel, L. & Woodgett, J. R. Lithium inhibits glycogen synthase kinase-3 activity and mimics wingless signalling in intact cells. Curr Biol 6, 1664–1668 (1996). 10.1016/s0960-9822(02)70790-2

42 Robles-Murguia, M., Hunt, L. C., Finkelstein, D., Fan, Y. & Demontis, F. Tissue-specific alteration of gene expression and function by RU486 and the GeneSwitch system. NPJ Aging Mech Dis 5, 6 (2019). 10.1038/s41514-019-0036-8

43 Küppers-Munther, B., et al. A new culturing strategy optimises Drosophila primary cell cultures for structural and functional analyses. Dev. Biol. 269, 459–478 (2004).

44 Dotti, C. G., Sullivan, C. A. & Banker, G. A. The establishment of polarity by hippocampal neurons in culture. J. Neurosci. 8, 1454–1468 (1988).

45 Lee, G., Neve, R. L. & Kosik, K. S. The microtubule binding domain of tau protein. Neuron 2, 1615–1624 (1989). 0896-6273(89)90050-0 [pii]

46 Hirokawa, N., Funakoshi, T., Sato-Harada, R. & Kanai, Y. Selective stabilization of tau in axons and microtubule-associated protein 2C in cell bodies and dendrites contributes to polarized localization of cytoskeletal proteins in mature neurons. J Cell Biol. 132, 667–679 (1996).

47 Grundke-Iqbal, I., et al. Abnormal phosphorylation of the microtubule-associated protein tau (tau) in Alzheimer cytoskeletal pathology. Proc Natl Acad Sci U S A 83, 4913–4917 (1986).

48 Schneider, C., Wicht, H., Enderich, J., Wegner, M. & Rohrer, H. Bone morphogenetic proteins are required in vivo for the generation of sympathetic neurons. Neuron 24, 861–870 (1999).

49 Doble, B. W. & Woodgett, J. R. GSK-3: tricks of the trade for a multi-tasking kinase. J Cell Sci 116, 1175–1186 (2003). 10.1242/jcs.00384

50 Wu, X., et al. Skin stem cells orchestrate directional migration by regulating microtubule-ACF7 connections through GSK3beta. Cell 144, 341–352 (2011).

51 Trivedi, N., Marsh, P., Goold, R. G., Wood-Kaczmar, A. & Gordon-Weeks, P. R. Glycogen synthase kinase-3beta phosphorylation of MAP1B at Ser1260 and Thr1265 is spatially restricted to growing axons. J Cell Sci 118, 993–1005 (2005). 10.1242/jcs.01697

52 Allen, J. J., et al. A semisynthetic epitope for kinase substrates. Nat Methods 4, 511–516 (2007). 10.1038/nmeth1048

53 Lauretti, E., Dincer, O. & Pratico, D. Glycogen synthase kinase-3 signaling in Alzheimer’s disease. Biochim Biophys Acta Mol Cell Res 1867, 118664 (2020). 10.1016/j.bbamcr.2020.118664

54 Tan, S., et al. Monoallelic loss-of-function variants in GSK3B lead to autism and developmental delay. Mol Psychiatry (2024). 10.1038/s41380-024-02806-z

55 Gobrecht, P., Leibinger, M., Andreadaki, A. & Fischer, D. Sustained GSK3 activity markedly facilitates nerve regeneration. Nat Commun 5, 4561 (2014). 10.1038/ncomms5561

56 Owen, R. & Gordon-Weeks, P. R. Inhibition of glycogen synthase kinase 3beta in sensory neurons in culture alters filopodia dynamics and microtubule distribution in growth cones. Mol Cell Neurosci 23, 626–637 (2003). 10.1016/s1044-7431(03)00095-2

57 Franco, B., et al. Shaggy, the Homolog of Glycogen Synthase Kinase 3, controls neuromuscular junction growth in Drosophila. J. Neurosci. 24, 6573–6577 (2004).

58 Cheng, Z., et al. Targeting glycogen synthase kinase-3beta for Alzheimer’s disease: Recent advances and future Prospects. Eur J Med Chem 265, 116065 (2024). 10.1016/j.ejmech.2023.116065

59 Hur, E. M., et al. GSK3 controls axon growth via CLASP-mediated regulation of growth cone microtubules. Genes Dev 25, 1968–1981 (2011). 25/18/1968 [pii] 10.1101/gad.17015911

60 Kumar, P., et al. GSK3beta phosphorylation modulates CLASP-microtubule association and lamella microtubule attachment. J Cell Biol 184, 895–908 (2009).

61 van Hummel, A., et al. No Overt Deficits in Aged Tau-Deficient C57Bl/6.Mapttm1(EGFP)Kit GFP Knockin Mice. PLoS One 11, e0163236 (2016). 10.1371/journal.pone.0163236

62 Ikegami, S., Harada, A. & Hirokawa, N. Muscle weakness, hyperactivity, and impairment in fear conditioning in tau-deficient mice. Neurosci Lett 279, 129–132 (2000). 10.1016/s0304-3940(99)00964-7

63 Goncalves, R. A., Wijesekara, N., Fraser, P. E. & De Felice, F. G. Behavioral Abnormalities in Knockout and Humanized Tau Mice. Front Endocrinol (Lausanne) 11, 124 (2020). 10.3389/fendo.2020.00124

64 Voelzmann, A., et al. Tau and spectraplakins promote synapse formation and maintenance through Jun kinase and neuronal trafficking. eLife 5, e14694 (2016).

65 Lee, J. & Kim, H. J. Normal Aging Induces Changes in the Brain and Neurodegeneration Progress: Review of the Structural, Biochemical, Metabolic, Cellular, and Molecular Changes. Front Aging Neurosci 14, 931536 (2022). 10.3389/fnagi.2022.931536

66 Okenve-Ramos, P., et al. Neuronal ageing is promoted by the decay of the microtubule cytoskeleton. PLoS Biol 22, e3002504 (2024). 10.1371/journal.pbio.3002504

67 Lee, S. J., et al. Age-related changes in glycogen synthase kinase 3beta (GSK3beta) immunoreactivity in the central nervous system of rats. Neurosci Lett 409, 134–139 (2006). 10.1016/j.neulet.2006.09.026

68 Souder, D. C. & Anderson, R. M. An expanding GSK3 network: implications for aging research. Geroscience 41, 369–382 (2019). 10.1007/s11357-019-00085-z

